# FoodMicrobionet v4: a large, integrated, open and transparent database for food bacterial communities

**DOI:** 10.1101/2022.01.19.476946

**Authors:** Eugenio Parente, Teresa Zotta, Annamaria Ricciardi

## Abstract

With the availability of high-throughput sequencing techniques our knowledge of the structure and dynamics of food microbial communities has made a quantum leap. However, this knowledge is dispersed in a large number of papers and hard data are only partly available through powerful on-line databases and tools such as QIITA, MGnify and the Integrated Microbial Next Generation Sequencing platform, whose annotation is not optimized for foods.

Here, we present the 4^th^ iteration of FoodMicrobionet, a database of the composition of bacterial microbial communities of foods and food environments. With 180 studies and 10,151 samples belonging to 8 major food groups FoodMicrobionet 4.1.2 is arguably the largest and best annotated database on food bacterial communities. This version includes 1,684 environmental samples and 8,467 food samples, belonging to 16 L1 categories and 196 L6 categories of the EFSA FoodEx2 classification and is approximately 4 times larger than previous version (3.1, https://doi.org/10.1016/j.ijfoodmicro.2019.108249).

Using data in FoodMicrobionet we confirm that taxonomic assignment at the genus level can be performed confidently for the majority of amplicon sequence variants using the most commonly used 16S RNA gene target regions (V1-V3, V3-V4, V4), with best results with higher quality sequences and longer fragment lengths, but that care should be exercised in confirming the assignment at species level.

Both FoodMicrobionet and related data and software conform to FAIR (findable, accessible, interoperable, reusable/reproducible) criteria for scientific data and software and are freely available on public repositories (GitHub, Mendeley data).

Even if FoodMicrobionet does not have the sophistication of QIITA, IMNGS and MGnify, we feel that this iteration, due to its size and diversity, provides a valuable asset for both the scientific community and industrial and regulatory stakeholders.

## 1. Introduction

In less than 15 years, next generation sequencing has added several layers of complexity to the study of food microbiomes. At the time of writing this article (January 2022) searches with the key words “microbiome” OR “microbiota” in Scopus and in Web of Science retrieved 118,416 and 127,424 hits, respectively. The number of papers using amplicon targeted next generation sequencing to study the structure of food bacterial communities has been rising exponentially since the first pioneering publications (Humblot and Guyot, 2009; Roh et al., 2010) (see Figure 1).

**Figure 1.**
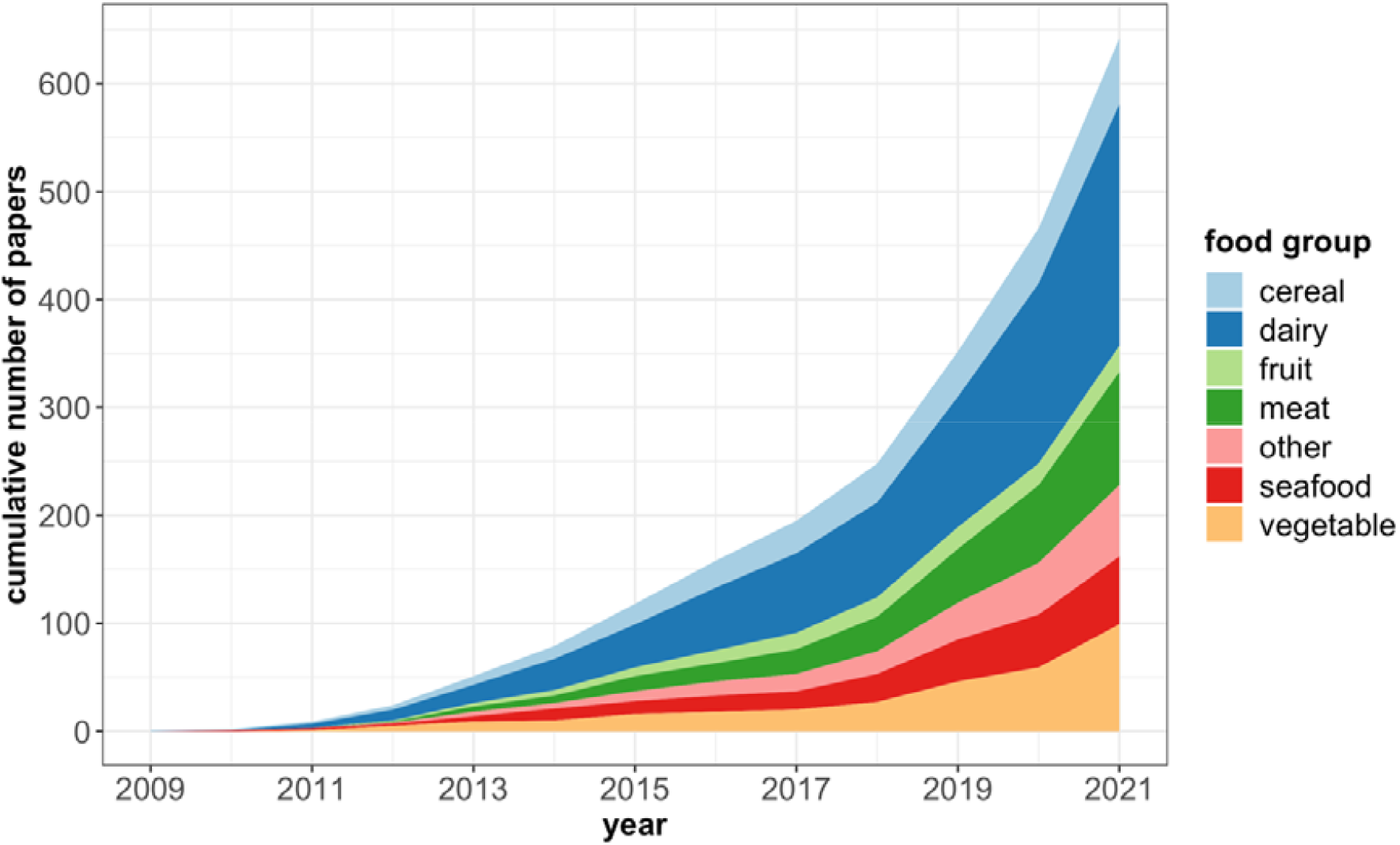
Cumulative number of papers using metataxonomic or metagenomic approaches for the study of food microbial communities. The data derive from a manually curated search on Web of Science™.

The evolution of -omic approaches for the study of the food microbiome and the methodological challenges in wet laboratory methods, sequencing and bioinformatic and statistical approaches have been reviewed multiple times (Bokulich et al., 2020; De Filippis et al., 2018; Pollock et al., 2018). While amplicon targeted approaches are still prevalent, the number of studies using metagenomic or multi-omics approaches is rapidly increasing with the decreasing cost of sequencing and the increase in power of sequencing platforms and computational tools (Yap et al., 2021).

Shotgun metagenomics offers significantly higher resolution (down to the strain and perhaps to single nucleotide polymorphism variants (Hildebrand, 2021) and paves the way to accurate source tracking for contamination (De Filippis et al., 2020) and to a new paradigm in food safety (Kovac, 2019). However, amplicon targeted approaches still have the power to describe, in a sensitive and cost-effective way, the structure of food microbial communities down to the genus level and possibly below (Callahan et al., 2016a; Johnson et al., 2019).

More than 640 papers describing the food microbiota have been published since 2009. Exploitation of this wealth of information for metastudies or for the development of descriptive or predictive tools requires FAIR (findable, accessible, interoperable, reusable/reproducible: Lamprecht et al., 2020; Wilkinson et al., 2016) data and software. Three large on line repositories on microbiome data have appeared in recent years: IMNGS (Lagkouvardos et al., 2016), QIITA (Gonzalez et al., 2018) and MGnify (Mitchell et al., 2019). The Integrated Microbial Next Generation Sequencing platform directly accesses 16S rRNA gene targeted next generation sequencing data in NCBI Short Read Archive and provides powerful tools for searching for taxa or sequences or for performing analysis of user-deposited 16S sequences (Lagkouvardos et al., 2016).

MGnify hosts metagenomic, metatranscriptomics and metataxonomic datasets which are either contributed by users or obtained from public repositories and offers powerful platforms for data analysis but does not allow the integration of data from different studies (Mitchell et al., 2019).

QIITA offers a powerful suite of tools for the analysis of sequences of public and private datasets, and allows the search and integration of data from different studies (Gonzalez et al., 2018).

At the time of writing of this paper (January 2022) MGnify included 3,696 public studies, and 325,323 public samples, but only 83 studies/2,805 samples on food biomes (https://www.ebi.ac.uk/metagenomics/) while QIITA included 620 public studies and 303,313 public samples but only a very limited number of public studies on food biomes (https://qiita.ucsd.edu/stats/). The data and interfaces for all these tools respond well to FAIR principles for scientific data (Wilkinson et al., 2016) but, unfortunately, the structure of the metadata for studies and samples is not optimized for foods.

Some years ago, we created a database for metataxonomic studies on food bacterial communities, FoodMicrobionet (Parente et al., 2016, 2019), whose main strength is the annotation system for studies and samples, based on the hierarchical classification of foods developed by the European Food Safety Authority, FoodEx 2 (E.F.S.A., 2015). Here we describe the structure and new features of the latest version of the database. In addition, we provide two proofs of concept on how the rich metadata structure of FoodMIcrobionet can be used to demonstrate the effect of target region on the resolution of taxonomic assignment of amplicon targeted metagenomics for food bacteria.

## 2. Materials and methods

### 2.1 Feeding data to FoodMicrobionet

The procedure used to add data to FoodMicrobionet has not changed since the last version (Parente et al., 2019) and it is represented schematically in Figure 2.

**Figure 2.**
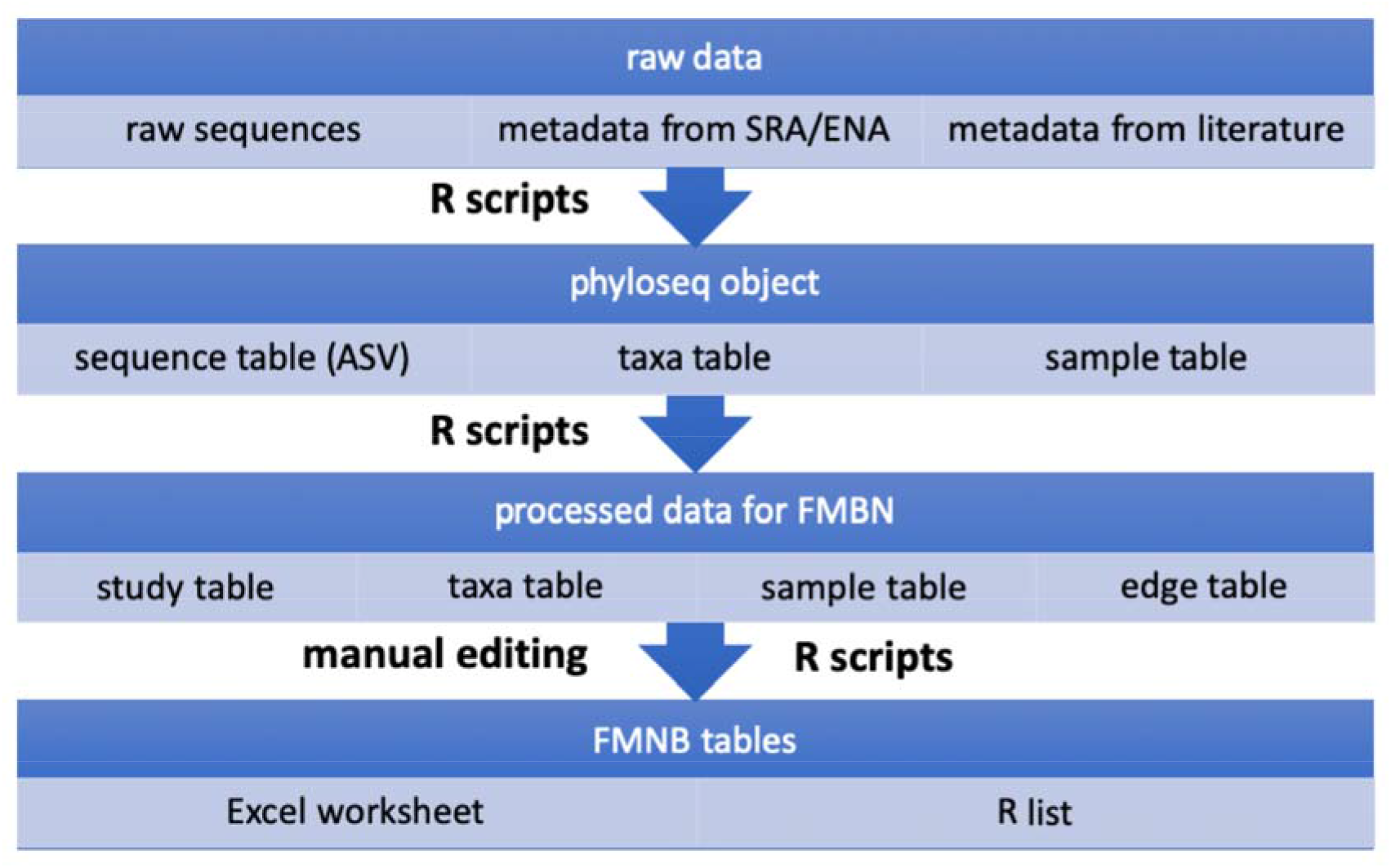
Schematic workflow showing how raw sequences and their metadata and additional information from the scientific literature are used to assemble data for FoodMicrobionet.

With the exception of 33 studies, which were originally provided by research partners as abundance tables for taxa and sample metadata tables (Parente et al., 2016), studies in FoodMicrobionet are added to the database by reprocessing raw sequence data from NCBI Short Read Archive (SRA), and by using metadata from SRA and from the scientific papers for annotation.

Processing of sequences is carried out in R (R Core Team, 2021) using a modified version of the Bioconductor pipeline for amplicon targeted sequence analysis, with DADA2 for Amplicon Sequence Variant (ASV) inference and SILVA v138.1 for taxonomic assignment (Callahan et al., 2016a; Callahan et al., 2016b). This results in the production of phyloseq (McMurdie and Holmes, 2013) objects which are processed using R scripts and imported in Microsoft Excel for further manual editing of study and sample metadata. Finally, a R script is used to process Excel tables and for quality control checks. All scripts are publicly available in the FoodMicrobionet GitHub repository (https://github.com/ep142/FoodMicrobionet).

### 2.2 The structure of FoodMicrobionet v4

The structure of the database is schematically shown in Figure 3.

**Figure 3.**
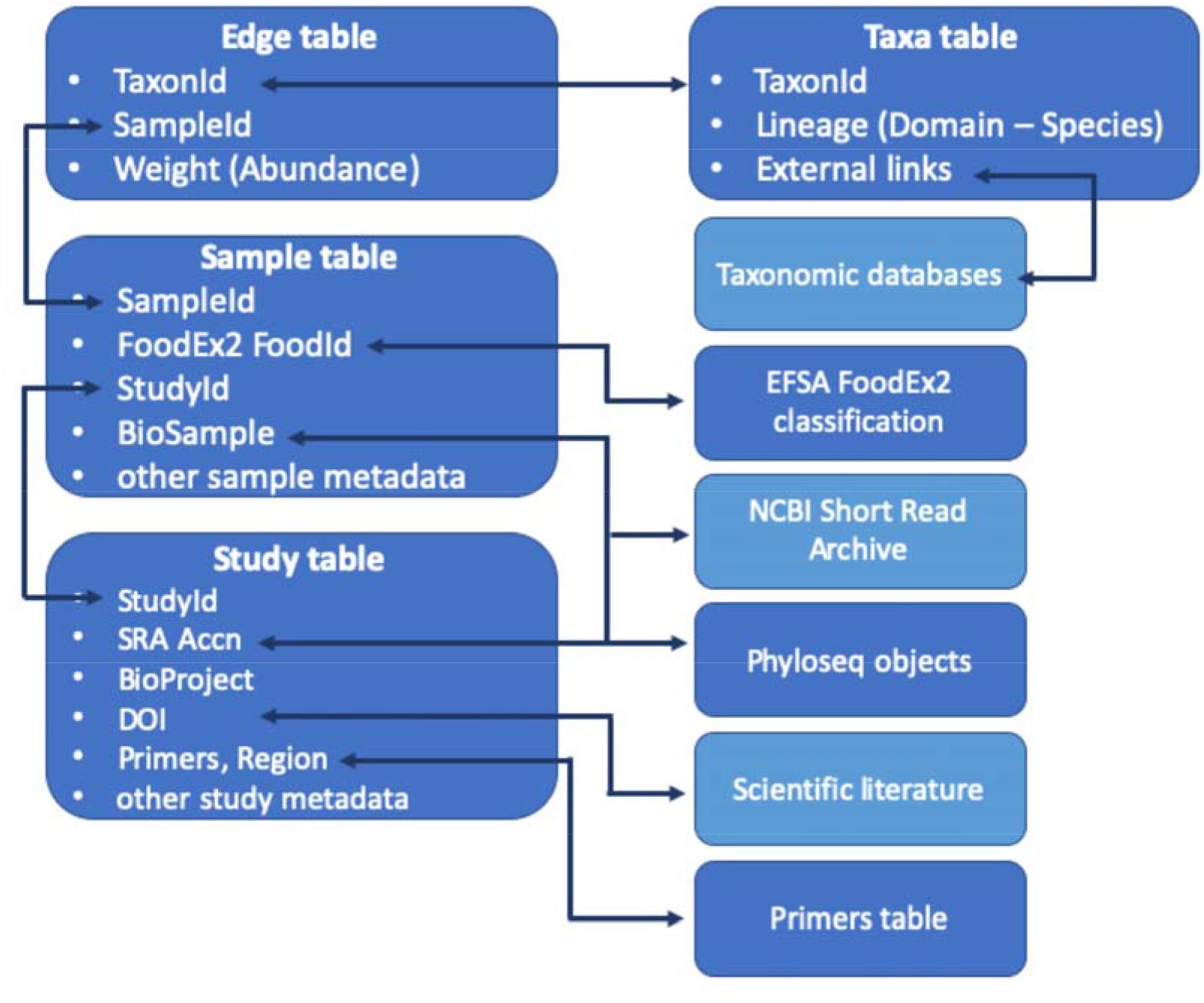
Schematic representation of the structure of FoodMicrobionet tables (Study, Primers, Samples, FoodEx2, Taxa, Edges) and their relationship with phyloseq (McMurdie and Holmes, 2013) objects and external databases.

Studies, samples and taxa have all a rich metadata structure. The structure of the tables is described in detail on GitHub (https://github.com/ep142/FoodMicrobionet/blob/master/FoodMicrobionet_tablespecs.md), on Mendeley data (https://data.mendeley.com/datasets/8fwwjpm79y/5) and in supplementary material.

Additions in version 4 include four fields which complete information for bioinformatic processing and one field for geolocation for studies, geolocation information for samples, and two further reference tables (primers and the FoodEx2 Exposure Hierarchy classification (E.F.S.A., 2015).

FoodMicrobionet is available either as R lists, which allow experienced programmers to run their searches and analyses in the most flexible and sensitive way, and as an interactive Shiny app (Parente et al., 2019). The latter requires minimum installation and configuration and allows users to perform searches using a large number of criteria, to perform aggregation of samples and taxa, to rapidly reach external resources using hyperlinks, to export data in a variety of formats, and to obtain and save graphs and tables (Parente et al., 2019).

### 2.3 Proof of concept 1: on the taxonomic resolution of amplicon targeted metagenomics for food bacterial communities

Using the metadata available in FoodMicrobionet we tabulated the frequency of taxonomic level assignments at the species, genus, family, order, class and phylum level for studies 34 to 180 (i.e. all studies for which sequence processing had been done using the procedure described in section 2.1). Graphs and tables were generated using a R script (ide_depth.R, available on GitHub, https://github.com/ep142/FoodMicrobionet/tree/master/the_real_thing/R_lists).

### 2.4 Proof of concept 2: using ASV for in depth analysis of taxonomic assignments

One of the major changes in FoodMicrobionet v4 is that ASV for each study can be directly accessed using the phyloseq objects created with the pipeline described in section 2.1. This in turn allows comparisons among ASV obtained in different studies using the same target region, and with reference sequences. To demonstrate this, we wrote a script which performed the following actions:

1. Search the database for two genera including pathogenic species (*Listeria, and Salmonella*) and identify the samples and studies in which they occur
2. Create graphs and tables showing their prevalence and abundance
3. Use study and sample accession numbers to retrieve ASV sequences from the phyloseq objects
4. Divide the sequences in groups (depending on the 16S RNA gene target region), carry out taxonomic assignment using RDP v18, and compare the sequences with reference sequences and outgroups extracted from the SILVA v138.1 reference database (two randomly extracted reference sequences for each species were used)
5. For each group, perform alignment and estimate Maximum Likelihood phylogenetic trees using the procedure described in Callahan et al. (2016b)
6. Annotate and plot phylogenetic trees using *treeio, tidytree* (Wang et al., 2020), and *ggtree* (Yu, 2020) R packages

## 3. Results and discussion

### 3.1 FoodMicrobionet facts and figures

FoodMicrobionet has grown significantly since its last iteration (Parente et al., 2019; Figure 4): the number of studies in version 4.1.2 has grown from 44 to 180, and the number of samples from 2,234 to 10,151.

**Figure 4.**
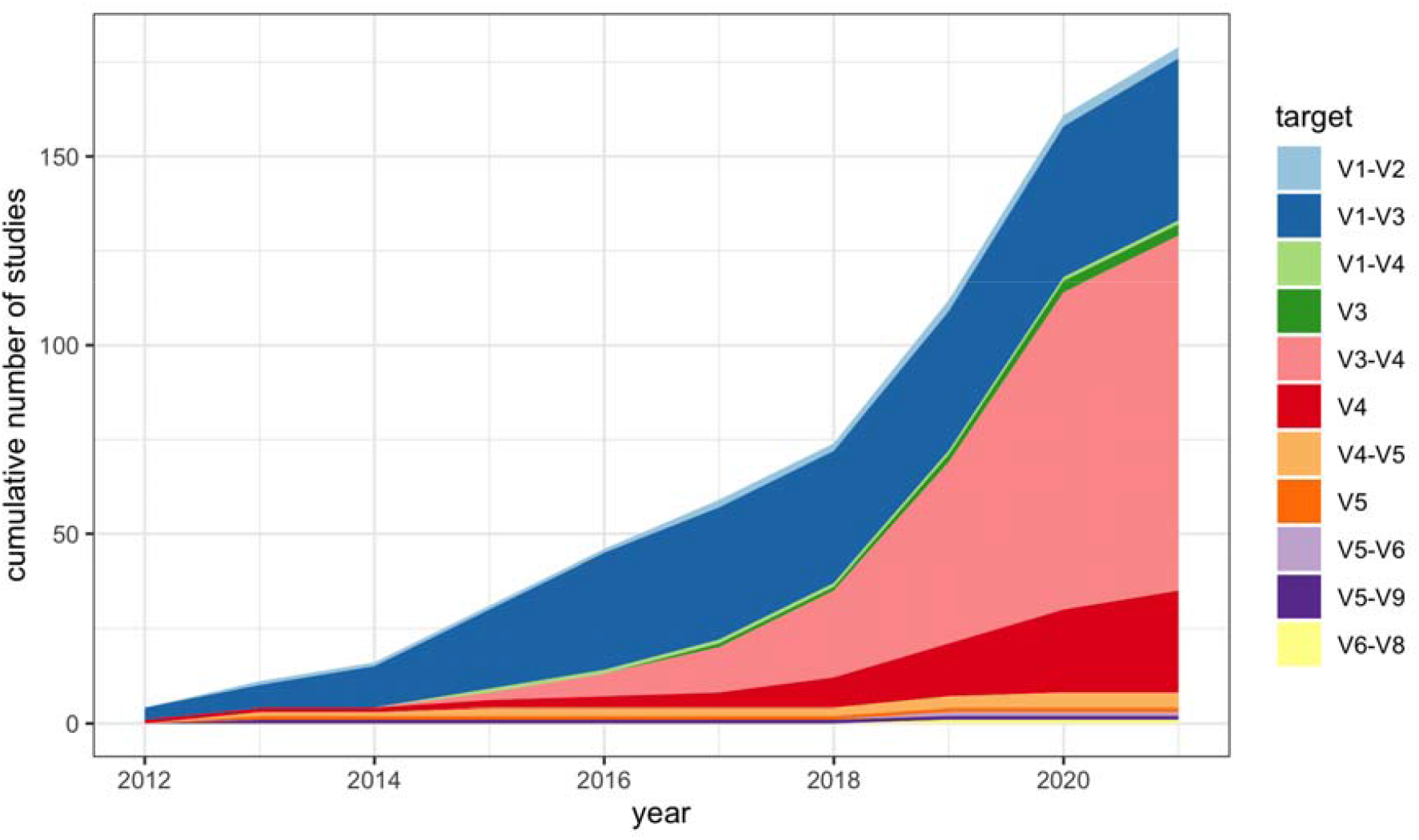
Cumulative distribution of studies in FoodMicrobionet 4.1.2, by year and by 16S rRNA gene target region.

The growth of FoodMicrobionet reflects the growth of published studies and the distribution of target gene regions reflects the shift in technology in Next Generation Sequencing, from the defunct Roche 454 platform (with most studies targeting the V1-V3 region) to Illumina platforms, with V4 and V3-V4 regions of the 16S rRNA gene as the most frequent targets (Figure 4, Supplementary Table 1).

The number of reads for sample after processing is also saved in the database (Supplementary Figure 1): 70% samples have >10^4^ reads after processing, but a significant number of samples in studies targeting the V4 region has >5×10^4^ reads. This makes the detection of rare components of the microbiota possible.

The variety of food groups and food types in FoodMicrobionet is also very large: samples in FMBN belong to 16 major food groups (L1 level of FoodEx2 exposure classification; Supplementary Table 2). Approximately 80% of samples belong to the first three categories (Milk and dairy products, Meat and meat products, Vegetables and vegetable products). Samples in FMBN are further classified using levels L4 and L6 of the FoodEx2 exposure classification, and additional fields (which allow to identify raw products, intermediates or finished products, the level of thermal treatment and the occurrence of spoilage and/or fermentation) allow a finer classification. Samples in FMBN belong to 109 L4 food groups and 197 L6 food groups. There are 163 individual food types, and, if further information on samples (nature, heat treatment, spoilage/fermentation) are used, there are 316 unique combinations (Parente et al., 2019). This number and variety of samples makes FoodMicrobionet the largest database of metataxonomic data on food bacterial communities, with significantly more samples compared to QIITA and MGnify. FoodMicrobionet stores samples from 51 countries, but the 90% of samples are from 14 countries (Supplementary Figure 2). This does not reflect the distribution of samples and studies in published studies but, rather, the distribution of those which are available in NCBI SRA.

There are currently 9,098 taxa in the taxa table of FoodMicrobionet. Taxa have a unique numeric and text identifier and may represent ASV identified at the species (4,497, 49.4% of taxa) or genus level (3,259, 35.8%) or above using the DADA2 assignTaxnomy() and assignSpecies() functions. The % of the taxa and sequences identified at the genus level or below varies by study, depending on the quality and length of sequences and on the gene target. Length of reads (in bp) in FoodMicrobionet studies varies between 150 and 610 bp (median 422).

FoodMicrobionet is fully connected to external databases: external links (as dynamically built Uniform Resource Locators) in the study, sample and taxa tables allow rapid access to external taxonomic databases (NCBI taxonomy, the List of Prokaryotic Names with Standing in Nomenclature and the Florilege database, Falentin et al., 2017 http://migale.jouy.inra.fr/Florilege/#&about) and to the scientific literature (via DOI). Using NCBI SRA Study accession number it is possible to access fine grained data on ASV sequence and abundance stored in the phyloseq objects obtained from processing the raw sequences (these are not public but are available on request).

### 3.2 Is FoodMicrobionet FAIR?

FoodMicrobionet data and software are free (both are under MIT licence https://opensource.org/licenses/MIT), open and highly reusable and support analysis protocols which are reproducible. We have done our best to conform as closely as possible to criteria for FAIR (findable, accessible, interoperable, reusable/reproducible) data and software sharing (Lamprecht et al., 2020; Wilkinson et al., 2016).

Both the database and the software are findable (using searches on Google, Mendeley data, or GitHub, for example) and deposited on permanent repositories (Mendeley data, GitHub) with unique identifiers. Since the database is not available on line except in the form of R lists or Excel files, data within FoodMicrobionet may not be directly findable by automated machine searches (and insofar they are not machine operable); however, the wealth of contextual metadata for all the main tables (studies, samples, taxa) makes it possible to devise precisely targeted searches.

FoodMicrobionet is accessible through the above-mentioned repositories and through our website, and we are confident that enough metadata are provided in these repositories to easily reach the resources. In terms of user accessibility (which is not a criterion in FAIR principles), we have done our best to make the resource accessible to both expert (the database in the form of a R list can be used for fine-tuned searches and analyses using R scripts) and moderately expert users. The latter can, with a minimum effort, download and install the R Shiny app, ShinyFMBN (Parente et al., 2019), which, once launched in R, allows easy access via an interactive and intuitive interface. A detailed manual for the app is available on Mendeley data (https://data.mendeley.com/datasets/8fwwjpm79y/4). Although the app could be easily deployed on RStudio Shiny apps server (https://www.shinyapps.io), we feel that this would make the use unnecessarily slow. FoodMicrobionet is fully interoperable. The sample classification is based on a robust hierarchical classification, FoodEx2, rather than on arbitrary keywords, and dynamic links are created in the studies, samples and taxa tables to reach a number of other databases. Conversely, other databases like Florilege might quite simply create new accessions using FoodMicrobionet and metadata in FMBN can, in principle, be used to populate QIITA and MGnify, by linking studies and samples via the SRA accession numbers.

Data and products of search results are highly reusable, for the same reasons. In addition, the objects exported by the app are in formats which are compatible with metataxonomic analysis pipelines (like phyloseq and ShinyPhyloseq: (McMurdie and Holmes, 2013, 2015); MicrobiomeAnalyst: (Chong et al., 2020; Dhariwal et al., 2017); CoNet: (Faust and Raes, 2016); graph visualization and analysis software like Cytoscape and iGraph: (Csardi et al., 2006; Shannon et al., 2003), microbial association network inference tools: (Kurtz et al., 2015; Peschel et al., 2021).

Use cases and example workflows have been illustrated in a previous work (Parente et al., 2019) and we have tried to demonstrate this approach in a series of proof-of-concept metastudies (Parente et al., 2020, 2021; Zotta et al., 2021). The code for generating graphs and statistical analyses is fully reproducible and reusable (https://data.mendeley.com/datasets/8fwwjpm79y/4; https://github.com/ep142/MAN_in_cheese) and allows to reproduce the figures and tables whenever a new version of FoodMicrobionet is published.

### 3.3 Proof of concept 1: on the taxonomic resolution of amplicon targeted metagenomics for food bacterial communities

The procedure for adding data to FMBN involves, starting from version 3 (studies 34 to 180) the use of a modification of the BioConductor workflow for microbiome data analysis (Callahan et al., 2016b). This procedure, which infers Amplicon Sequence Variants using DADA2, has performed well in benchmarking and can be used to compare data across multiple studies (Callahan et al., 2016a. Callahan et al., 2017). ASV inference is increasingly been used in the study of food bacterial communities (55% of studies 160 to 180 in FoodMicrobionet originally used DADA2; see Supplementary Table 3 for references). The ability to perform taxonomic assignment to the lowest possible level (species) is clearly important in food microbial ecology, because different members a given genus may have different roles in foods (beneficial, detrimental, pathogenic); this is for example the case for *Clostridium, Bacillus, Staphylococcus, Corynebacterium*, and many other genera of food related bacteria. Different variable regions of the 16S RNA gene have been historically used as targets for amplicon targeted metagenomics, and FoodMicrobionet provides a comprehensive sampling of the targets used (Figure 4).

The median and 90° percentile values for the frequency of taxonomic assignment at the genus level or below in FMBN studies from 34 to 180 is shown in Supplementary Table 4. The weighted (by sequence abundance, Figure 5) and unweighted (Supplementary Figure 3) frequency of assignment at the genus level or below varied between studies, even within a given target region (Figure 5). This was apparently related to sequence length and target region, and a clear relationship was found with at least one indicator of average within-study sequence quality. In fact, for paired end sequences for the longest target regions, merging of paired ends was not always possible due to bad quality of sequences toward the 5’ end (data not shown) and this prevented the overlap of forward and reverse sequences. For these sequences, the BioConductor workflow used in FoodMicrobionet allows taxonomic assignment down to the genus level, but not down to the species level (Callahan et al., 2016b).

**Figure 5.**
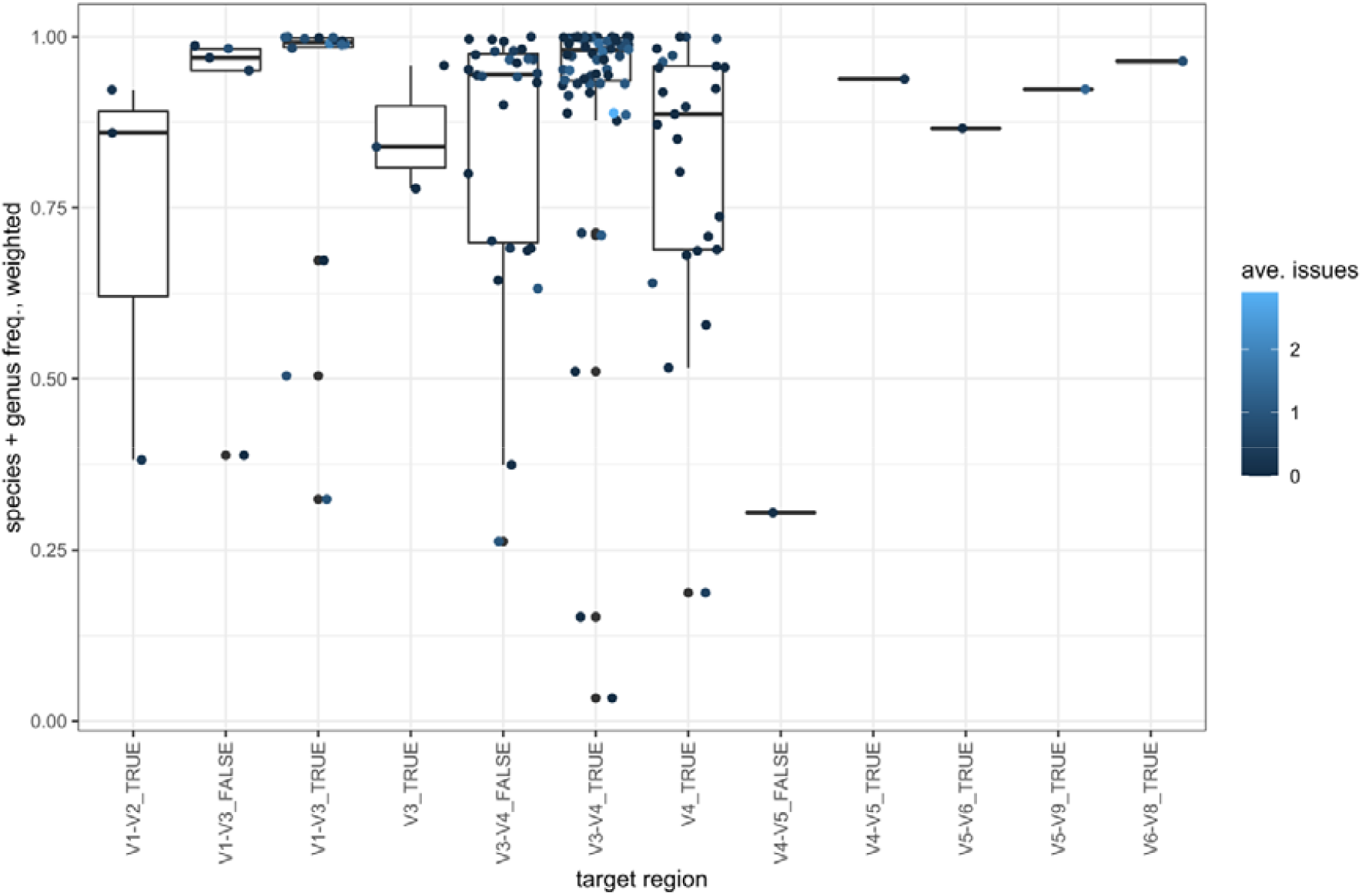
Box and jitter plots showing the weighted (by sequence relative abundance) distribution of frequencies of taxonomic assignments at the genus level or below in FoodMicrobionet studies 34 to 180. The average values for the number of issues encountered during bioinformatic processing (high sequence losses during filtering or chimera removal, low number of final sequences, low diversity) is also shown.

However, no clear relationship was found with the other indicator of sequence quality provided by FoodMicrobionet, i.e. the average number of issues during bioinformatic processing, see table specifications in Supplementary material). Overall, the median value of the frequency of taxonomic assignment at the genus level or below ranged from 0.640 and 0.898, with the lowest values for the shortest regions and studies with the worse sequence quality. However, when the number of sequences for each ASV is taken into account these figures may change significantly, and median values for taxonomic assignment at the genus level or below as high as 0.98 (overlapping V3-V4 region) or 0.99 (overlapping V1-V3 region) can be obtained (Figure 5). Shorter regions still provide a reasonably good performance (with weighted median frequencies of genus assignments of 0.73 and 0.80 for V3 and V4 respectively). However, species assignments were much less frequent, with median values for weighted frequencies ranging from 0.025 to 0.381 (Supplementary Table 5). Median weighted values for species level assignment were 0.38 and 0.30 for regions V1-V3 and V3-V4, respectively, but as low as 0.21 for V4, a frequently used target in large recent studies. Differences in the frequencies of taxonomic assignment were also observed for different phyla. This is illustrated for the four most abundant phyla in FoodMicrobionet (*Firmicutes, Proteobacteria, Actinobacterota, Bacteroidota*; SILVA taxonomy is used for higher taxa) in Supplementary Figures 4 and 5. For some targets the ability to perform taxonomic assignment at the genus level or below was clearly lower and/or more variable.

These results are in good agreement with a recent study which compared taxonomic assignment for different target regions within the 16S RNA gene (Johnson et al., 2019). The possibility of assigning taxonomy down to the species level was found to differ among regions, with species level assignments are significantly less frequent for shorter regions compared to the full 16S RNA gene, which can now be sequenced using 3^rd^ generation High Throughput Sequencing platforms, like PacBio and Oxford Nanopore (Johnson et al., 2019). In addition, differences in the ability to perform taxonomic assignment at the genus and species level for different phyla may also explain why the weighted and unweighted frequencies of identification differ: abundant sequences often belong to taxa which are well represented in taxonomic reference databases (data not shown).

In FoodMicrobionet, target region and composition of the microbiota of the study are clearly not independent. Given the metadata structure in FoodMicrobionet one could, at least in principle, compare the taxonomic assignment for different food groups (which might, in turn, reflect differences in microbial community composition), but this might result in too many different combinations. However, at least for the target regions for which a large number of studies is available (V3-V4, V4, and, to a lesser extent, V1-V3) we feel that the results offer a wide enough coverage of food groups and clearly indicate that for *Actinobacterota* and *Bacteroidota* the pipeline used in FoodMicrobionet may offer a lower degree of success in taxonomic assignment compared to *Proteobacteria* and *Firmicutes*.

### 3.4 Proof of concept 2: using Amplicon Sequence Variants for in depth analysis of taxonomic assignments

To further demonstrate the need to exert caution in taxonomic assignment down to the species level with relatively short targets we developed a second proof of concept. First, we searched the database for samples containing members of genera *Listeria* and *Salmonella*. While metagenomic approaches, due to their high resolution, are certainly more useful than metataxonomic approaches in food safety studies (Cocolin et al., 2018; Jagadesaan et al., 2019; Kovac, 2019), the latter may still have value for studying the microbial ecology of food borne pathogens. However, this requires taxonomic assignment to a level which is low enough to discriminate species, or species groups, which are relevant for human or animal health. The distribution of the abundance of members of the genera *Listeria* and *Salmonella* in L1 food categories of the EFSA FoodxEx2 classification is shown in Figure 6.

**Figure 6.**
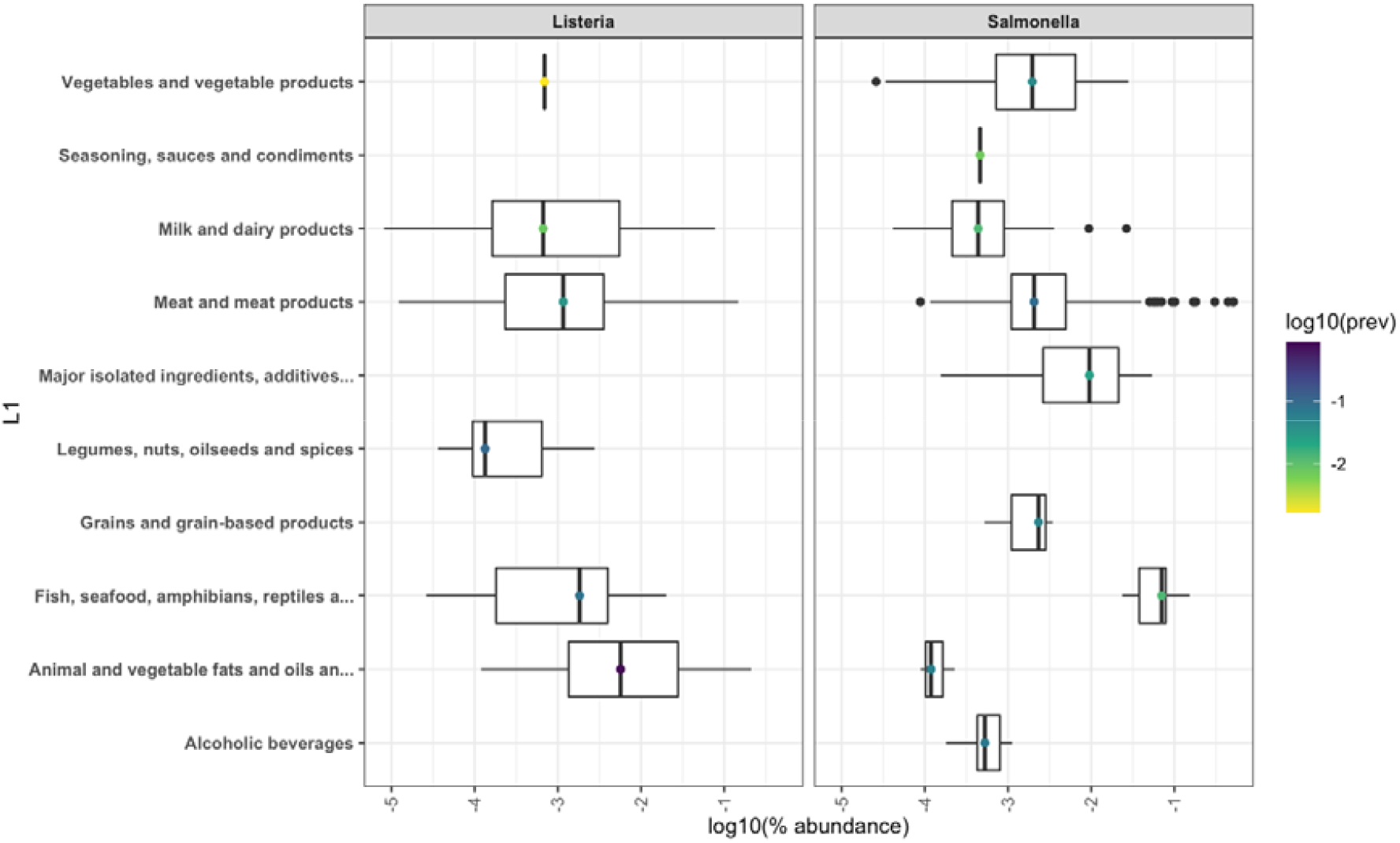
Distribution of the abundance of six genera including pathogenic bacteria in FoodMicrobionet samples. The L1 level of food classification of the EFSA FoodEx2 classification is shown. Prevalence is shown as a colour scale. Environmental samples were excluded from the analysis.

With some exceptions in which abundance was >0.1% and as high as 0.5% of total sequences (Supplementary Table 6), these genera occur with a low prevalence and abundance (typically <0.01%) and would normally be discarded by abundance and prevalence filters which are normally applied when processing microbiome data (Callahan et al., 2016b). The low abundance and prevalence and the occurrence of some genera in unexpected environments (*like Salmonella in alcoholic beverages*) may rise the doubt that their detection is due to contamination or to errors in sequence processing or taxonomic assignment. Unfortunately, although it is well known that contamination may severely affect the results of microbiome analysis, especially in low biomass samples (Dahlberg et al., 2019; Davis et al., 2018; Pollock et al., 2018), the use of blanks and control and the application of statistical procedures for the removal of contamination (Davis et al., 2018) is very rare. Moreover, even if mock communities may assist in benchmarking the bioinformatic processing of sequences in metataxonomic studies (Bokulich et al., 2020; Pollock et al., 2018) they, too, are very rarely used in food microbial ecology studies.

While providing conclusive results on the occurrence of contamination or on the quality of bioinformatic processing is impossible, the accessibility of ASV in FoodMicrobionet using study and sample accession numbers provides an opportunity for checking the quality of the taxonomic assignment and carry out direct comparison with reference sequences.

We therefore re-identified all sequences belonging to *Listeria* and *Salmonella* using the RDP v18 trainset reference database. The results are shown in Supplementary Figure 6. While for *Listeria* the two databases resulted in matching identifications, for *Salmonella*, a high proportion of sequences were assigned to other genera (*Enterobacter, Citrobacter*) using RDP. Differences in taxonomic assignment due to the use of different reference databases are not surprising (Ramakodi, 2022; Werner et al., 2012) even if SILVA and RDP often produce matching assignments (Smith et al., 2020).

Finally, we generated phylogenetic trees for sequences grouped by region, and included reference sequences from the SILVA v138.1 taxonomic reference database, including outgroups (*Brochothrix thermosphacta* for *Listeria* and *Citrobacter freundii* for *Salmonella*). Reference sequences for *Enterobacter cloacae* were also included for *Salmonella*. Results for overlapping paired sequences for region V3-V4 are shown in Figure 7, while combined results for the V3-V4 and V4 region are shown in Supplementary Figure 7. Results with other regions were similar (data not shown). Classification obtained by probabilistic assignment (as in the naïve Bayesian classifier, Wang, 2007) and phylogenetic tree inferences are based on different approaches and are not easy to compare.

**Figure 7.**
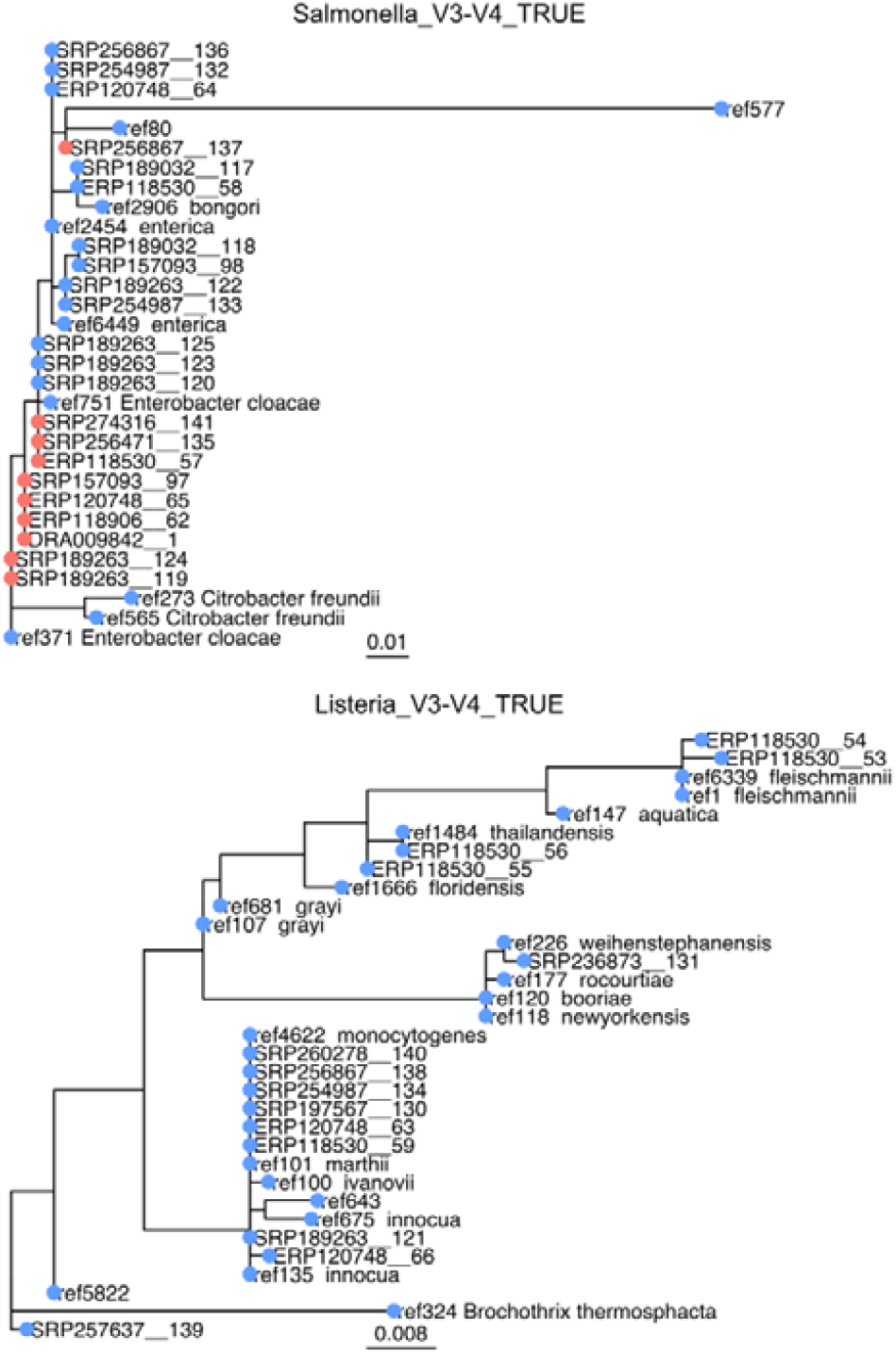
Maximum likelihood phylogenetic trees for amplicon sequence variants (ASVs) for overlapping paired end sequences for region V3-V4 of the 16S RNA gene identified as *Salmonella* or *Listeria*. ASVs are identified by the accession number of the study to which they belong and by a random progressive integer. Reference sequences extracted from SILVA v138.1 taxonomic reference database are also included. Colored dots indicate sequences for which taxonomic assignment with SILVA v138.1 and RDP trainset 18 matched (blue) or not (red).

However, for *Listeria*, reference sequences for different species grouped in several clades (with slight differences in grouping for different regions). Due to small phylogenetic distance within each clade, with reference sequences differentiated by a very small number of nucleotide changes, it is not surprising that species assignment by the naïve Bayesian classifier was often not successful. A single ASV (SRP257637_139) did not group with *Listeria* reference sequences. As to *Salmonella*, the majority of ASVs grouped with *Salmonella* reference sequences, even when taxonomic assignment at the species level with RDP was different. However, at least for the V3-V4 region, one reference sequence for *Enterobacter cloacae* clustered with *Salmonella*. This may explain differences in assignment at the genus level with the two databases.

It is well known that accuracy of taxonomic assignment with the naïve Bayesian classifier may vary for different regions (Wang, 2007) and that sequences in reference databases may have erroneous taxonomic annotations (Pollock et al., 2018). However, we feel that our results confirms that the ability to perform taxonomic assignments varies with different regions of the 16S RNA gene (Johnson et al., 2019), and when using short fragments, even when a taxonomic assignment at the species level is obtained, one should be wary of the results.

Although one may question the value of taxonomic assignment at the species level, especially for the shortest reads (Callahan et al., 2016a; Edgar, 2018; Johnson et al., 2019; Meola et al., 2019; Pollock et al., 2018), due to the detailed information provided for both studies and samples (gene target and region, the number of issues observed during processing of raw sequences) and to the possibility of accessing to the processed sequences (ASV) in phyloseq objects, users of FoodMicrobionet can make informed decision on how and when taxonomic units should be combined at a level higher than the species, an operation which is easily performed with the ShinyFMBN app (Parente et al., 2019).

## 4. Conclusions

Even if FoodMicrobionet does not have the sophistication of QIITA, IMNGS and Mgnify, we feel that this iteration, due to its size and diversity, provides a significant resource for both the scientific community and industrial and regulatory stakeholders. Scientists can access and use a variety of stand-alone or online software tools and ShinyFMBN to compare their own results with literature results, carry out metastudies to answer a variety of scientific questions, build reproducible analysis workflows, get quantitative data on the ecology/distribution of bacteria of interest, use the database as an entry point for further searches in other databases. The size of this version, which includes >9×10^5^ taxon/sample relationships, might even allow the machine learning approaches to predict contamination patterns of food. The ability to rapidly retrieve information on prevalence/abundance of taxon in different foods and on the structure of microbial communities in different food types may be useful to both the industry and regulatory agencies. Information on the distribution of beneficial genera and, to a lesser extent, species may find use for regulatory purposes (for example to facilitate studies on the distribution of beneficial microorganisms to evaluate their inclusion in the Qualified Presumption of Safety). The fine-grained data on the structure of microbial communities for a large variety of raw materials, foods, food environments may be useful for both process and product development purposes to identify spoilage or contamination patterns, or for the design of microbiome-based starters. We are committed to keep adding data to FoodMicrobionet, but the openness and transparency of its software and documentation allows any interested party to create new versions of the database or to significantly improve its structure and functionality.

## Supporting information

Table specifications

## CrediT author statement

**Eugenio Parente:** Conceptualization, Methodology, Software, Writing-Original draft preparation. **Annamaria Ricciardi** Data curation, Writing – Reviewing and Editing. **Teresa Zotta:** Data curation, Writing – Reviewing and Editing.

## Data statement

The database and related scripts and apps are available on GitHub (https://github.com/ep142/FoodMicrobionet) and on Mendeley data (https://data.mendeley.com/datasets/8fwwjpm79y/6). Phyloseq objects are available upon request.

## Acknowledgements

This research did not receive any specific grant from funding agencies in the public, commercial, or not-for-profit sectors.

