## Supplementary material for "FoodMicrobionet v4: a large, integrated, open and transparent database for food bacterial communities": Table specifications: FMBNtablespecs.html

FoodMicrobionet Tables specifications


### FoodMicrobionet Tables specifications

###### 14 January 2022

### **FoodMicrobionet 4.1: Structure of edges and nodes tables**.

This document describes the structure of data tables for FoodMicrobionet (FMBN) version 4.1.

The structure of the database was changed and is now more coherent with that of a relational database, to facilitate extraction of data. It now includes:

1. **studies table** : data on the studies included in FMBN. Includes info on the target (16S DNA or cDNA or both, region) sequencing platform, bioinformatic pipeline, sequence accession number, number of samples, bibliographic information; linked to the other tables by studyID
2. **primers table**: information on primers
3. **samples table** (sample nodes): data on the samples included in FMBN, see below; linked to the study table by studyId and to edge table by sampleId
4. **FoodEx2 table** : for convenience only; this table includes all codes from Exposure hierarchy from revision 2
5. **taxa table** (OTU nodes, formerly lineages table): data on taxa included in FMBN, see below; linked to the edge table by taxonId
6. **edges table** (edges, the source is a sample, the target a taxon, formerly edge table): data on sample - taxon edges, with abundances; linked to the study table by studyId, the sample table by sampleId, the taxon table by taxonId

The structure of the database is illustrated in Figure 1. Phyloseq objects are connected to studies via studyId but not included in the main database.

**Figure 1**. Structure and relationships in FoodMicrobionet. FoodMicrobionet tables are in dark blue. External tables connected through links are in turquoise.

The emphasis is now on data extraction and processing using a range of R Shiny apps, and most of the new features were designed to facilitate processing with other software packages (Microsoft Excel, Gephi, Cytoscape, the CoNet App of Cytoscape, the bipartite package and the phyloseq package of R). To learn more on FMBN visit our web site at: http://www.foodmicrobionet.org.

All scripts are available at https://github.com/ep142/FoodMicrobionet.

#### **The studies table**.

This table lists all relevant information for the study. The table has the following fields (each study is a record)

1. **study** : a numerical (integer) id for the study
2. **studyId** : an ID for the study (in the format STx, where x is an integer with the study number); this is assigned as studies are added to FMBN (see samples table)
3. **FMBN\_version** : the version of FMBN in which the study appeared for the first time (unless stated otherwise, the study will appear in all further versions)
4. **target** : the gene target(s) used for amplification
5. **region** : the region of the 16S RNA gene target used for amplification
6. **platform** : the sequencing platform
7. **read\_length\_bp** : the average read length
8. **seq\_center** : the sequencing centre
9. **bioinf\_software** : the bioinformatics software/pipeline used for the analysis
10. **OTU\_picking** : the OTU picking method
11. **assign\_tax\_method** : the taxonomy assignment method
12. **tax\_database** : the taxonomic database used for annotation
13. **seq\_accn** : the accession number for the SRA study, if available, empty otherwise (will become NA in R)
14. **seq\_accn\_link** : the hyperlink to the run selector for the SRA study, if available, otherwise will point to a page in the FMBN website
15. **bioproject** : SRA bioproject id if available
16. **samples** : the number of samples included in FMBN
17. **food\_group** : the food group (for tabulation purposes)
18. **short\_descr** : a short description of the study including FoodId (see samples table)
19. **DOI** : the DOI of the study, if published, "unpublished" otherwise
20. **DOI\_link** : the DOI hyperlink
21. **ref\_short** : an abbreviated reference to the published paper, if available
22. **year** : publication year; if missing, the year the experiment was carried out
23. **ref\_complete** : a complete reference to the paper describing the study, "unpublished" otherwise
24. **corr\_author** : corresponding author surname
25. **corr\_author\_mail** : corresponding author e-mail address
26. **geoloc** : a list of the countries in which the samples were obtained; see samples table
27. **primer\_f** : forward primer used for amplification (see primers table)
28. **primer\_r** : reverse primer used for amplification (see primers table)
29. **overlapping** : logical flag indicating if during processing paired end sequences overlapped correctly (if not a 10N spacer was added between forward and reverse, see documentation for DADA2)
30. **paired\_end** : logical flag indicating if paired end sequences were available^1^

#### **The primers table**.

The **primers** table includes some information on the primers (when available). This can be useful if a user wants to repeat the processing of raw sequences. The following fields are included

1. **primer** : the name of the primer (must be unique)
2. **sequence** : the sequence of the primer without adapters and spacers
3. **region** : the 16S region
4. **start** : start nucleotide
5. **length** : length in bp (calculated from sequence)
6. **note:** notes

#### **The samples table**.

The **samples** table includes metadata for samples/nodes. The current version of FMBN includes the following fields:

1. **studyId** : an ID for the study (in the format STx, where x is an integer with the study number), link to the same field in the studies table
2. **sampleId** : an integer with the number of the sample (samples are given a progressive number when added to FMBN), link to the same field in the edges table
3. **label\_1** : an ID for the sample (in the format SAy, where y is an integer with the number of the sample),
4. **label\_2** : character label for the sample, as given when the sample was added to FMBN. For compatibility with older versions of FMBN, the original sample label (i.e. the label as it appears in the original study or in SRA RunInfo tables under SampleName, whichever was available when the sample was added to FMBN) may have a prefix in the format xxx\_
5. **label\_3** : this column contains the node display label (used as node display label in interactive visualisations produced with Gephi); built by combining label\_2 and foodId
6. **llabel** : incorporates information on foodId and food status (see below) in a 10-character format (5 characters for the foo did, 3 for food status, see below for abbreviations, 1 for target, either d for 16S RNA gene or r for 16S RNA).^2^
7. **s\_type** : sample type: "Sample" for nodes corresponding to food samples, and "Environment" for samples from food environments.
8. **n\_reads** : number of reads after clean-up, empty if not available
9. **n\_reads2** : n\_reads if available, otherwise a dummy value (3227, the median of the values of n\_reads for samples for which this information is available in FMBN v 2) to be used for samples (food or environment) if n\_reads is not available; this value is used to rebuild OTU tables with absolute frequencies starting from the edge and samples tables
10. **n\_issues** : number of issues during sequence processing (i.e. high number of chimeras/bimeras, high loss during quality filtering, low number of features, low number of sequences after processing); set to -1 for studies 1:33 to avoid issues with column parsing when importing with readr functions
11. **Ontology fields, samples/environment** : the hierarchical terms and IDs are derived from the food classification and description system FoodEx 2 (revision 2) adopted by the European Food Safety Authority (EFSA, 2015). The terms used in FoodMicrobionet are derived from the exposure hierarchy. For food environments they are set to the same values of the food processed in that environment.^3^

- **foodId** : the code from the FoodEx 2 classification
- **description** : a description of the sample, including at least terms from L6
- **L1** : the top level in the hierarchy
- **L4** : level 4 in the hierarchy
- **L6** : level 6 in the hierarchy.

12. **Food status fields** : these columns describe the status of the food sample (empty environmental samples). They are inferred from what is known (from the original paper, from information provided by the contributor) on a sample^4^

- **nature** : **Raw** (raw material or ingredient), **Intermediate** , **Finished** (including the products during storage). Shorthands: r, i, f; 0 for environmental samples or when this information is not available.
- **process** : **None** (raw materials and foods other than those covered by the other options); **Mild** (foods with no physical lethal treatment; includes foods with added preservatives, mild heat treatments and fermentation); **Type2** (foods submitted at least to a lethal treatment resulting potentially in a 6D decrease in *L. monocytogenes*, but whose treatment is lower than that of type 1 foods; in these foods the heat treatment is between 70°C for 2 min and 90°C for 10 min); **Type1** (foods submitted at least to a lethal treatment resulting potentially in a 6D decrease in *C. botulinum* type E spores, approximately 90°C for 10 min or equivalent)^5^. Shorthands: n, m, 2, 1; 0 for environmental samples or when this information is not available.
- **spoilage**^6^: **NA** (no information available), **Unspoiled** (all fresh or processed, non spoiled foods); **Fermented** (all unspoiled fermented foods), **Spoiled** (all non-fermented foods which are spoiled or are at the end of shelf-life), **Fermented+spoiled** (fermented foods which are also spoiled or at the end of shelf-life). Shorthands: 0 for environmental samples or when this information is not available.

13. **target1** : the gene used as a target
14. **target2** : the region used as a target
15. **links to NCBI SRA database**:^7^

- **biosample**: SRA biosample, if available, otherwise empty
- **SRA\_sample**: SRA sample, if available, otherwise empty
- **SRA\_run**: SRA run, if available, otherwise empty

16. **geolocation information** : geolocation information derived from SRA metadata tables, when possible or from the published papers

- **geo\_loc\_country** : the country where the sample was collected
- **geo\_loc\_continent** : the continent where the sample was collected
- **lat\_lon** : the coordinates of the place where the sample was collected

Future iterations of this table may contain further metadata on environmental variables (pH, a_W_, etc.) as measured or estimated values.

#### **The FoodEx2 exposure table**.

This table is included for reference purposes, since a few fields are already included in the Samples table. However, users may conceivably want to use level of aggregation (L2, L3, L5) which are not included in FoodMicrobionet or pool FoodId codes under a code of higher hierarchical level.

1. **Code** : the unique identifier for a food or food category, matches FoodId in the samples table (unique)
2. **L1** : the level 1 in the classification, matches L1 in the samples table
3. **L2** : the level 1 in the classification
4. **L3** : the level 1 in the classification
5. **L4** : the level 1 in the classification, matches L4 in the Samples table
6. **L5** : the level 1 in the classification
7. **L6** : the level 1 in the classification, unique, matches L6 in the Samples table

#### **The taxa table**.

The **taxa** table includes information on the lineages of taxa included in FMBN and outlinks to external resources (NCBI taxonomy, List of Prokaryotic Names with Standing in Nomenclature, Florilege. It is linked to the edges table by the taxonId field.

1. **taxonId** : an integer with the number of the taxon (taxa are given a progressive number when added to FMBN as a result of the addition of a sample-taxon relationship)
2. **label** : a label for the taxon; must be unique; corresponds to the binomial for the species (if available) or to a higher taxon (genus, family, etc. + \_unidentified) when the OTU was not identified at the species level
3. **lineage fields** : the order is not used in this version; when the information is not available the field is left empty

- **domain**
- **phylum**
- **class**
- **order**
- **family**
- **genus**
- **species**

4. **id\_L6** the taxonomic assignment including the lineage in the format^8^: Root:k\_\_Bacteria\_\_Firmicutes:c\_\_Bacilli:f\_\_Lactobacillaceae:g\_\_Lactobacillus:s\_\_Lactobacillusdelbrueckii
5. **taxonomy** : a lineage built for compatibility with the CoNet app of Cytoscape
6. **idelevel** : the taxonomic level of identification (domain, …, species)
7. **NCBI\_outlink** : outlink to NCBI taxonomy
8. **BacterioNet\_outlink** : outlink to LPSN
9. **Flori\_habitat** : outlink to the Florilege database (taxon lives in habitat tab)
10. **Flori\_pheno** : outlink to the Florilege database (taxon exhibits phenotype tab)
11. **Flori\_use** : outlink to the Florilege database (taxon studied for use tab)

#### **The edges table**.

The **edges table** includes a list of all relationships between taxa and samples (food or food environment). The fields included in the current version of FoodMicrobionet are:

1. **sampleId** (Food/food environment sample): an integer with the number of the sample (samples are given a progressive number when added to FMBN), link to the same field in the samples table
2. **taxonId** : an integer with the number of the taxon, link to the same field of the taxa table
3. **weight** : the OTU abundance in the sample: for (FMBN) historical reasons this is in %^9^ but is converted in frequency (0-1) or absolute abundance during filtering and selection in the ShinyFMBN app.

#### **References**.

- EFSA (2015). The food classification and description system FoodEx 2 (revision 2). Supporting publications EFSA Supporting publication 2015:EN-804. http://www.efsa.europa.eu/en/supporting/pub/215e.htm
- Parente, E., Cocolin, L., De Filippis, F., Zotta, T., Ferrocino, I., O'Sullivan, O., Neviani, E., De Angelis, M., Cotter, P. D., Ercolini D. (2016). FoodMicrobionet: A database for the visualisation and exploration of food bacterial communities based on network analysis. *International Journal of Food Microbiology*, *219*, 28–37. http://doi.org/10.1016/j.ijfoodmicro.2015.12.001

*-* Parente, E., De Filippis, F., Ercolini, D., Ricciardi, A., Zotta, T., 2019. Advancing integration of data on food microbiome studies: FoodMicrobionet 3.1, a major upgrade of the FoodMicrobionet database. *International Journal of Food Microbiology*, 305, 108249. https://doi.org/10.1016/j.ijfoodmicro.2019.108249

---

1. This is not necessarily true for all sequences obtained by Illumina, because in some cases only forward sequences were deposited in SRA↩︎
2. this is one of the fields used for filtering and labelling (in graphs)↩︎
3. there is an issue with food samples obtained during processing of, for example, a hard, ripened cheese. A cheese curd is quite obviously a completely different environment compared to a mature hard cheese. This issue is dealt with by the "Nature" field; when the correct classification is not found in FoodEx or information are partially missing the closest relative is used.↩︎
4. in the future we may add info on intrinsic factors (pH, aW) or extrinsic factors (temperature used during storage/ripening, atmosphere); however, this information is (alas!) often missing and may need to be inferred by knowledge on the process; as an alternative one may add facets from FoodEx2, see table 22 and 25 of the document. However, this may be difficult because sample description in published studies is often incomplete.↩︎
5. this is applicable to ready to eat foods. Both type1 and type2 may include fermented foods which have been submitted to a heat treatment ACMSF, 2007. Report on safe cooking of burgers. Retrieved from http://www.food.gov.uk/multimedia/pdfs/acmsfburgers0807.pdf.↩︎
6. includes foods at the end of shelf life, even if they do not show clear spoilage.↩︎
7. in the first iteration of version 3 these are available only for samples belonging to studies form 34 on; the other studies will be fixed later.↩︎
8. It is important to understand that the original taxonomic lineage obtained when directly analysing sequences from SRA/ENA or from contributors is manually edited for coherence. Contributors have submitted in the past OTU abundance tables with lineages in different formats and obtained from different version of taxonomic reference databases; as a result, the same taxon often had a different lineage, especially for *incertae sedis* taxa. When a new lineage is added several steps are carried out: lineages pointing to the same (at the genus, family, class, phylum level) taxon are merged; the lineage is checked against SILVA classification, bad taxa (taxa missing one level of classification) are fixed and a few genera are edited to allow linking to external database (*Corynebacterium\_1* becomes *Corynebacterium*). When any of the taxonomic terms (k\_\_; p\_\_, c\_\_, etc.) the corresponding column is left blank.↩︎
9. this was needed to set correctly node size in Gephi; proportions or number of sequences can be obtained simply by dividing by 100 or by using information in the Samples table.↩︎
