## Supplementary tables and figures for "FoodMicrobionet v4: a large, integrated, open and transparent database for food bacterial communities"

Scuola di Scienze Agrarie, Forestali, Alimentari ed Ambientali, Università degli Studi della Basilicata, Potenza, Italy

**Supplementary Tables and Figures**

**Supplementary Table 1.** Distribution of samples in FoodMicrobionet, by target 16S rRNA gene region and sequencing platform

| region | 454 GS | Illumina | Ion Torrent | Sum |
| --- | --- | --- | --- | --- |
| V1-V2 | 1 | 1 | 1 | 3 |
| V1-V3 | 33 | 11 | 0 | 44 |
| V1-V4 | 1 | 0 | 0 | 1 |
| V3 | 0 | 3 | 0 | 3 |
| V3-V4 | 1 | 93 | 0 | 94 |
| V4 | 2 | 22 | 3 | 27 |
| V4-V5 | 2 | 2 | 0 | 4 |
| V5 | 1 | 0 | 0 | 1 |
| V5-V6 | 0 | 1 | 0 | 1 |
| V5-V9 | 1 | 0 | 0 | 1 |
| V6-V8 | 0 | 1 | 0 | 1 |
| Sum | 42 | 134 | 4 | 180 |

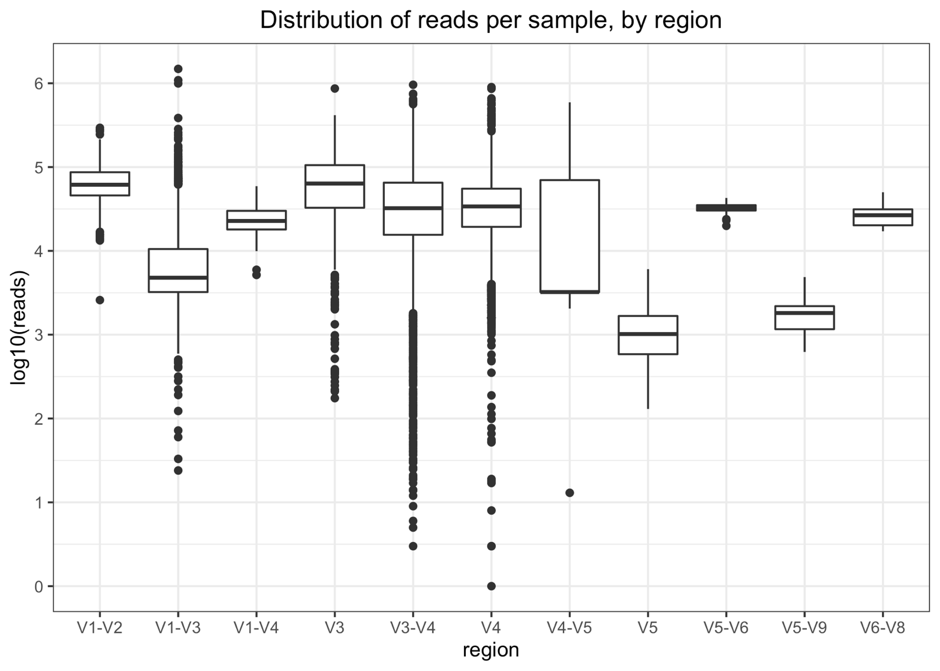

**Supplementary Figure 1**. Distribution of reads per sample in FoodMicrobionet, by target 16S rRNA gene region, after processing.

**Supplementary Table 2.** Number of samples (n, food or environmental samples) belonging to different categories of the Level 1 in FoodEx2 classification. The categories are arranged in decreasing order of size (in terms of number of samples) and the proportion (prop) and the cumulative proportion of samples (cumprop) is shown.

| L1 | n | prop | cumprop |
| --- | --- | --- | --- |
| Milk and dairy products | 4918 | 0.4845 | 0.4845 |
| Meat and meat products | 2641 | 0.2602 | 0.7447 |
| Vegetables and vegetable products | 604 | 0.0595 | 0.8042 |
| Fruit and fruit products | 525 | 0.0517 | 0.8559 |
| Fish, seafood, amphibians, reptiles and invertebrates | 493 | 0.0486 | 0.9044 |
| Major isolated ingredients, additives, flavours, baking and processing aids | 280 | 0.0276 | 0.9320 |
| Seasoning, sauces and condiments | 138 | 0.0136 | 0.9456 |
| Composite dishes | 115 | 0.0113 | 0.9570 |
| Alcoholic beverages | 114 | 0.0112 | 0.9682 |
| Legumes, nuts, oilseeds and spices | 81 | 0.0080 | 0.9762 |
| Grains and grain-based products | 66 | 0.0065 | 0.9827 |
| Eggs and egg products | 56 | 0.0055 | 0.9882 |
| Animal and vegetable fats and oils and primary derivatives thereof | 53 | 0.0052 | 0.9934 |
| Sugar and similar, confectionery and water-based sweet desserts | 42 | 0.0041 | 0.9975 |
| Starchy roots or tubers and products thereof, sugar plants | 17 | 0.0017 | 0.9992 |
| Water and water-based beverages | 8 | 0.0008 | 1.0000 |

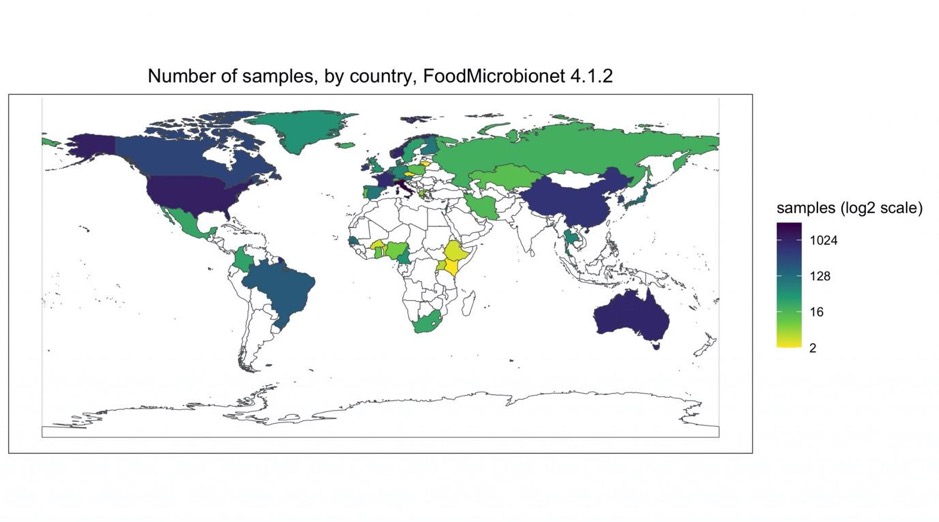

**Supplementary Figure 2**. The geography of FoodMicrobionet 4.1.2: number of food and food environment samples, by country.

**Supplementary Table 3**. Studies included in FoodMicrobionet.

| Study Id | FMBN version | Reference |
| --- | --- | --- |
| ST1 | 0_6_2 | Guidone, A., Zotta, T., Matera, A., Ricciardi, A., De Filippis, F., Ercolini, D., Parente, E., 2015. The microbiota of high-moisture mozzarella cheese produced with different acidification methods. Int. J. Food Microbiol. 216: 9-17. |
| ST2 | 0_6_2 | Parente, E., Guidone, A., Matera, A., De Filippis, F., Mauriello, G., Ricciardi, A., 2016. Microbial community dynamics in thermophilic undefined milk starter cultures. Int. J. Food Microbiol. 217: 59-67. |
| ST3 | 0_6_2 | De Filippis, F., La Storia, A., Stellato, G., Gatti, M., Ercolini, D., 2014. A selected core microbiome drives the early stages of three popular Italian cheese manufactures. PLoS One 9 (2): e89680. |
| ST4 | 0_6_2 | Ercolini, D., De Filippis, F., La Storia, F., Iacono, M., 2012. "Remake" by High-Throughput Sequencing of the microbiota involved in the production of water buffalo Mozzarella cheese. Appl. Environ. Microbiol. 78: 8142-8145. |
| ST5 | 0_6_2 | Marsh, A. J., O'Sullivan, O., Hill, C., Ross, R. P., Cotter, P. D., 2013. Sequencing-based analysis of the bacterial and fungal composition of kefir grains and milks from multiple sources. PLoS One 8 (7): e69371. |
| ST6 | 0_6_2 | De Pasquale, I., Calasso, M., Mancini, L., Ercolini, D., La Storia, A., De Angelis, M., Di Cagno R., Gobbetti, M., 2014. Causal relationship between microbial ecology dynamics and proteolysis during manufacture and ripening of Canestrato Pugliese PDO Cheese. Appl. Environ. Microbiol. 80: 4085-4094. |
| ST7 | 0_6_2 | De Pasquale, I., Di Cagno, R., Buchin, S., De Angelis, M., Gobbetti, M., 2014. Microbial ecology dynamics reveal a succession in the core microbiota involved in the ripening of pasta filata Caciocavallo Pugliese Cheese. Appl. Environ. Microbiol. 80: 6243-6255. |
| ST8 | 0_6_2 | Dolci, P., De Filippis, F., La Storia, A., Ercolini, D., Cocolin, L., 2014. rRNA-based monitoring of the microbiota involved in Fontina PDO cheese production in relation to different stages of cow lactation. Int. J. Food Microbiol. 185: 127-135. |
| ST9 | 0_6_2 | Alessandria, V.,Ferrocino, I., De Filippis, F., Fontana, M., Rantsiou, K., Ercolini, D., Cocolin, L., 2016. Microbiota of an Italian Grana like cheese during manufacture and ripening unraveled by 16S rRNA-based approaches. Appl. Environ. Microbiol. 82: 3988-3995. |
| ST10 | 1_0_3 | De Filippis, F., Genovese,A., Ferranti, P., Gilbert, J. A., Ercolini, D. 2016. Metatranscriptomics reveals temperature-driven functional changes in microbiome impacting cheese maturation rate. Sci. Rep. 6:21871. |
| ST11 | 1_0_3 | De Filippis, F. La Storia, A., Villani, F., Ercolini, D., 2013. Exploring the sources of bacterial spoilers in beefsteaks by culture-independent High-Throughput Sequencing. PLoS One 8 (7): e70222. |
| ST12 | 1_0_3 | Greppi, A., Ferrocino, I., La Storia, A., Rantsiou, K., Ercolini, D., Cocolin, L., 2015. Monitoring of the microbiota of fermented sausages by culture Independent rRNA-based approaches. Int. J. Food Microbiol. 212: 67-75. |
| ST13 | 1_0_3 | Ferrocino, I., Greppi, A., La Storia, A., Rantsiou, K., Ercolini, D., Cocolin, L., 2015. Impact of nisin-activated packaging on microbiota of beef burgers during storage. Appl. Environ. Microbiol. 82: 549-559. |
| ST14 | 1_0_3 | Cocolin, L., Alessandria, V., Botta, C., Gorra, R.,De Filippis, F., Ercolini, D., Rantsiou, K., 2013. NaOH-debittering induces changes in bacterial ecology during table olives fermentation. PLoS One 8 (7): e69074. |
| ST15 | 1_0_3 | Ercolini et al., unpublished data |
| ST16 | 1_0_3 | Rizzello, C. G., Cavoski, I., Turk, J., Ercolini, D., Nionelli, L., Pontonio, E., De Angelis, M., De Filippis, F., Gobbetti, M., Di Cagno, R., 2015. Organic cultivation of Triticum turgidum subsp. durum Is reflected in the flour-sourdough fermentation-bread axis. Appl. Environ. Microbiol. 81: 3192–3204. |
| ST17 | 1_0_3 | Ercolini, D. Pontonio, E., De Filippis, F., Minervini, F., La Storia, A., Gobbetti, M., Di Cagno, R., 2013. Microbial ecology dynamics during rye and wheat sourdough preparation. Appl. Environ. Microbiol. 79: 7827-7836. |
| ST18 | 1_1 | Stellato, G., De Filippis, F., La Storia, A., Ercolini, D., 2015. Coexistence of lactic acid bacteria and potential spoilage microbiota in a dairy processing environment. Appl. Environ. Microbiol. 81: 7893-7904. |
| ST19 | 1_1 | Pothakos, V., Stellato, G., Ercolini, D., Devlieghere, F,. 2015. Processing environment and ingredients are both sources of Leuconostoc gelidum, which emerges as a major spoiler in ready-to-eat meals. Appl. Environ. Microbiol. 81: 3529-3541. |
| ST20 | 1_1 | Stellato, G., La Storia, A., Cirillo, T., Ercolini, D., 2015. Bacterial biogeographical patterns in a cooking center for hospital foodservice. Int. J. Food Microbiol. 193: 99-108. |
| ST21 | 1_1 | Stellato, G., La Storia, A., De Filippis, F., Borriello, G., Villani, F., Ercolini, D., 2016. Overlap of spoilage microbiota between meat and meat processing environment in small-scale vs large-scale retail distribution. Appl. Environ. Microbiol., 82:4045–4054 |
| ST22 | 1_1 | Calasso, M., Ercolini, D., Mancini, L., Stellato, G., 2016. Relationships among house, rind and core microbiotas during manufacture of traditional Italian cheeses at the same dairy plant. Food Microbiol. 54: 11-26. |
| ST23 | 1_1 | Bassi, D., Puglisi, E., Cocconcelli, P. S., 2015. Understanding the bacterial communities of hard cheese with blowing defect. Food Microbiol. 52: 106–118. |
| ST24 | 1_1 | Rebecchi, A., Pisacane, V., Miragoli, F., Polka, J., Falasconi, I., Morelli, L., Puglisi, E., 2015. High-throughput assessment of bacterial ecology in hog, cow and ovine casings used in sausages production. Int. J.l Food Microbiol. 212: 49-59. |
| ST25 | 1_1 | O'Sullivan, D. J., Cotter, P. D., O'Sullivan, O., Giblin, L.,McSweeney, P. L. H., Sheehan, J. J., 2015. Temporal and spatial differences in microbial composition during the manufacture of a continental-type cheese. Appl. Environ. Microbiol. 81: 2525-2533. |
| ST26 | 2 | De Angelis, M., Campanella, D., Cosmai, L., Summo, C., Rizzello, C. G., Caponio, F., 2015. Microbiota and metabolome of un-started and started Greek-type fermentation of Bella di Cerignola table olives. Food Microbiol. 52: 18-30. |
| ST27 | 2 | Minervini, F., Lattanzi, A., De Angelis, M., Celano, G., Gobbetti, M., 2015. House microbiotas as sources of lactic acid bacteria and yeasts in traditional Italian sourdoughs. Food Microbiol. 52:66-76. |
| ST28 | 2 | Ceuppens, S., Delbeke, S., De Coninck, D., Boussemaere, J., Boon, N., Uyttendaele, M., 2015. Characterization of the bacterial community naturally present on commercially grown basil Leaves: evaluation of sample preparation prior to culture-independent techniques. Int. J. Environ. Res. Publ. Health 12: 10171-10197. |
| ST29 | 2 | Hultman, J., Rahkila, R., Ali, J., Rousu, J., Björkroth, K. J., 2015. Meat processing plant microbiome and contamination patterns of cold-tolerant bacteria causing food safety and spoilage risks in the manufacture of vacuum-packaged cooked sausages. Appl. Environ. Microbiol. 81: 7088-7097. |
| ST30 | 2 | Leff, J .W., Fierer, N., 2013. Bacterial communities associated with the surfaces of fresh fruits and vegetables. PLoS One 8 (3): e59310. |
| ST31 | 2 | Chaillou, S., Chaulot-Talmon, A., Caekebeke, H., Cardinal, M., Christieans, S., Denis, C., Desmonts, M. H., Dousset, X.,Feurer, C., Hamon, E., Joffraud, J.-J., La Carbona, S., Leroi, F., Leroy, S., Lorre, S., Macé, S., Pilet, M.-F., Prévost, H., Rivollier, M., Roux, D., Talon, R., Zagorec, M., Champomier-Vergès, M.-C., 2015, Origin and ecological selection of core and food-specific bacterial communities associated with meat and seafood spoilage. The ISME J. 9: 1105–1118. |
| ST32 | 2 | Quigley, L., O'Sullivan, O., Beresford, T. P., Ross, R. P., Fitzgerald, G. F., Cotter, P. D., 2012. High-throughput sequencing for detection of subpopulations of bacteria not previously associated with artisanal cheeses. Appl. Environ.l Microbiol. 78: 5717-5723. |
| ST33 | 2 | Quigley, l., O'Sullivan, D. J., Daly, D., O’Sullivan, O., Burdikova, Z., Vana, R., Beresford, T. P., Ross, R. P., Fitzgerald, G. F., McSweeney, P. L. H., Giblin, L., Sheehan, J. J.,  Cotter. P. D., 2016. Thermus and the pink discoloration defect in cheese. mSystems 1(3): e00023-16. |
| ST34 | 3_1 | Sattin, E., Andreani, N. A., Carraro, L., Fasolato, L. , Balzan, S., Novelli, E., Squartini, A., Telatin, A., Simionati, B., Cardazzo, B., 2016. Microbial dynamics during shelf-life of industrial ricotta cheese and identification of a Bacillus strain as a cause of a pink discolouration. Food Microbiol. 57 (August): 8–15. |
| ST35 | 3_1 | Sattin,E., Andreani, N. A., Carraro, L., Lucchini, R., Fasolato, L., Telatin, A., Balzan, S., Novelli, E., Simionati,B., Cardazzo, B., 2016. A multi-omics approach to evaluate the quality of milk whey used in ricotta cheese production. Front. Microbiol. 7: 1272. |
| ST36 | 3_1 | Minervini, F., Conte, A., Del Nobile, M. A., Gobbetti, M., De Angelis, M., 2017. Dietary fibers and protective lactobacilli drive burrata cheese microbiome. Appl. Environ. Microbiol. 83 (21): e01494-17 |
| ST37 | 3_1 | Doyle, C. .J, Gleeson, D., O'Toole, P. W., Cotter, P. D., 2017. High-throughput metataxonomic characterization of the raw milk microbiota Identifies changes reflecting lactation stage and storage conditions. Int. J. Food Microbiol. 255 (August): 1–6. |
| ST38 | 3_1 | Porcellato, D. Aspholm, M., Borghild Skeie, S., Monshaugen, M., Brendehaug, J., Mellegård, H., 2018. Microbial diversity of consumption milk during processing and storage. Int. J. Food Microbiol. 266 (February): 21–30. |
| ST39 | 3_1 | Frétin, M., Martin, B., Rifa, E., Verdier-Metz, I., Pomiès, D., Ferlay, A., Montel, M.-C., Delbès, C. 2018. Bacterial community assembly from cow teat skin to ripened cheeses Is Influenced by grazing systems. Sci. Rep. 8 (1): 200. |
| ST40 | 3_1 | Marino, M., Innocente, N., Maifreni, M., Mounier, J., Cobo-Díaz, J. F., Coton, E., Carraro, L., Cardazzo, B., 2017. Diversity within Italian cheesemaking brine-associated bacterial communities evidenced by massive parallel 16S rRNA gene tag eequencing. Front.Microbiol. 8: 2119. |
| ST41 | 3_1 | Dugat-Bony, E., Garnier, L., Denonfoux, J., Ferreira,S., Sarthou, A.-S., Bonnarme, P., Irlinger, F., 2016. “Highlighting the microbial diversity of 12 French cheese varieties. Int. J. Food Microbiol. 238: 265–73. |
| ST42 | 3_1 | Walsh, A. M., Crispie, F., Kilcawley, K., O'Sullivan, O., G O'Sullivan, M., Claesson, M. J., Cotter, P. D., 2016. Microbial succession and flavor production in the fermented dairy beverage Kefir. mSystems 1 (5): e00052–16. |
| ST43 | 3_1 | Guzzon, R., Carafa, I., Tuohy, K., Cervantes, G., Vernetti, L., Barmaz, A., Larcher,R., Franciosi, E., 2017. Exploring the microbiota of the red-brown defect in smear-ripened cheese by 454-pyrosequencing and Its prevention using different cleaning systems. Food Microbiol. 62: 160-168. |
| ST44 | 3_1 | Giello, M., La Storia, A., Masucci, F., Di Francia, A., Ercolini, D., Villani, F., 2017. Dynamics of bacterial communities during manufacture and ripening of traditional Caciocavallo of Castelfranco cheese in relation to cows' feeding. Food Microbiol. 63:170-177. |
| ST45 | 3_2 | Salazar, J. K., Carstens, C. K., Ramachandran,P., Shazer, A. G., Narula, S. S., Reed, E., Ottesen, A., Schill, K. M., 2018. Metagenomics of pasteurized and unpasteurized Gouda cheese using targeted 16S rDNA sequencing. BMC Microbiol. 18 (1): 189. |
| ST46 | 3_2 | Skeie, S. B., Håland, M., Thorsen, I. M.,Narvhus, J., Porcellato, D., 2019. Bulk tank raw milk microbiota differs within and between farms: a moving goalpost challenging quality control. J. Dairy Sci. 102 (3): 1959–1971. |
| ST47 | 3_2 | Cremonesi, P. Ceccarani, C., Curone, G., Severgnini, M., Pollera, C., Bronzo, V., Riva,F., Addis, M. F., FilipeI, J., Amadori, M., Trevisi, E., Vigo, D., Moroni, P., Castiglioni. B., 2018. Milk microbiome diversity and bacterial group prevalence in a comparison between healthy Holstein Friesian and Rendena cows.PLoS ONE 13 (10): e0205054. |
| ST48 | 3_2 | Marino, M., Dubsky de Wittenau, G., Saccà, E., Cattonaro, F., Spadotto, A., Innocente, N., Radovic, S., Piasentier, E., Marroni, F., 2019. Metagenomic profiles of different types of Italian high moisture Mozzarella cheese. Food Microbiol. 79 (June): 123–131. |
| ST49 | 3_2 | Kamimura, B. A., De Filippis, F., Sant'Ana, A. S., Ercolini, D., 2019. Large-scale mapping of microbial diversity in artisanal Brazilian cheeses. Food Microbiol. 80 (June): 40-49. |
| ST50 | 3_2 | Quijada, N. M., De Filippis, F.,Sanz, J. J., Del Camino García-Fernández, M., Rodríguez-Lázaro, D., Ercolini, D., Hernández, M., 2018. Different Lactobacillus populations dominate in 'Chorizo De León' manufacturing performed in different production plants. Food Microbiol. 70 (April): 94-102. |
| ST51 | 3_2 | Ferrocino, I., Bellio, A., Romano,A., Macori, G., Rantsiou, K., Decastelli, L., Cocolin, L., 2017. RNA-based amplicon sequencing highlights the impact of the processing technology on microbiota composition of artisanal and industrial PDO Lard d'Arnad. Appl. Environ. Microbiol. 83:e00983-17 |
| ST52 | 3_2 | Nieminen, T. T., Koskinen, K., Laine, P.,Hultman, J., Säde,E., Paulin, L., Paloranta,A., Johansson,P ., Bjorkroth, J., Auvinen, P., 2012 Comparison of microbial communities in marinated and unmarinated broiler meat by metagenomics. Int. J. Food Microbiol. 157 (2): 142–149. |
| ST53 | 3_2 | Fougy, L., Desmonts,M.-H., Coeuret,G., Fassel, C., Hamon,E., Hezard, B., Champomier-Vergès, M. C., Chaillou, S., 2016. Reducing salt in raw pork sausages increases spoilage and correlates with reduced bacterial diversity. Appl. Environ. Microbiol. 82 (13): 3928–3939. |
| ST54 | 3_2 | Botta, C., Ferrocino, I., Cavallero, M. C., Riva, S., Giordano, M., Cocolin, L., 2018. Potentially active spoilage bacteria community during the storage of vacuum packaged beefsteaks treated with aqueous ozone and electrolyzed water. Int. J. Food Microbiol. 266 (February): 337-345. |
| ST55 | 3_2 | Giello, M., La Storia, A., De Filippis, F., Ercolini, D., Villani, F., 2017. Impact of Lactobacillus curvatus 54M16 on microbiota composition and growth of Listeria monocytogenes in fermented sausages. Food Microbiol. 72 (November): 1-15. |
| ST56 | 3_2 | Cardinali, F., Milanović,V., Osimani,A., Aquilanti, L., Taccari, M., Cristiana Garofalo, Polverigiani, S., Clementi, F., Francios, E., Tuohy, K., Mercuri,M.L., Altissimi, M., Haouet, M. N., 2018. Microbial dynamics of model Fabriano-like fermented sausages as affected by starter cultures, nitrates and nitrites. Int. J. Food Microbiol. 278 (August): 61-72. |
| ST57 | 3_2 | Raimondi, S., Luciani, S., Sirangelo, T. M., Amaretti, A., Leonardi, A., Ulrici, A., Foca, G., D'Auria, G., Moya, A., Zuliani, V., Seiberth, T. M., Søltoft-Jenseni, J., Rossi, M., 2019. Microbiota of sliced cooked ham packaged in modified atmosphere throughout the shelf life. Int. J. Food Microbiol. 289 (January): 200–208. |
| ST58 | 3_2 | Juárez-Castelán, C., García-Cano,I., Escobar-Zepeda, A., Azaola-Espinosa, A., Álvarez-Cisneros, Y., Ponce-Alquicira, E., 2019. Evaluation of the bacterial diversity of Spanish-type Chorizo during the ripening process using High-Throughput Sequencing and physicochemical characterization. Meat Sci. 150 (April): 7–13. |
| ST59 | 3_2 | Lauritsen, C. V., Kjeldgaard,J., Ingmer, H., Bisgaard, M., Christensen, H., 2019. Microbiota encompassing putative spoilage bacteria in retail packaged broiler meat and commercial broiler abattoir. Int. J. Food Microbiol. 300 (July): 14–21. |
| ST60 | 3_2 | Chen, X., Zhang, Y., Yang, X., Hopkins, D. L., Zhu, L., Dong, P., Liang, R., Luo, X., 2019. Shelf-life and microbial community dynamics of super-chilled beef imported from Australia to China. Food Res. Int. 120 (June): 784–792. |
| ST61 | 3_2 | Zotta, T., Parente, E., Ianniello, R. G., De Filippis, F., Ricciardi, A., 2019. Dynamics of bacterial communities and interaction networks in thawed fish fillets during chilled storage in air. Int.l J. Food Microbiol. 293 (January):102-113. |
| ST62 | 3_2 | Pimentel, T., Marcelino, J., Ricardo, F., Soares, A. M. V. M., Calado, R., 2017. Bacterial communities 16S rDNA fingerprinting as a potential tracing tool for cultured Seabass Dicentrarchus labrax. Sci. Rep. 7 (1): 11862. |
| ST63 | 3_2 | Jääskeläinen, E., Jakobsen, L. M. A., Hultman, J., Eggers, N., Bertram, H. C., Björkroth, J., 2019. Metabolomics and bacterial diversity of packaged yellowfin tuna (Thunnus albacares) and salmon (Salmo salar) show fish species-specific spoilage development during chilled storage. Int. J. Food Microbiol. 293 (March): 44–52. |
| ST64 | 3_2 | Vieira, D. A. P., Cabral, L., Noronha, M. F., Júnior, G. V. L., Sant'Ana, A. S., 2019. Microbiota of eggs revealed by 16S rRNA-based sequencing from raw materials produced by different suppliers to chilled pasteurized liquid products. Food Control 96 (February): 194–204. |
| ST65 | 3_2 | Margot, H., Stephan, R., Tasara, T., 2016. Mungo bean sprout microbiome and changes associated with culture based enrichment protocols used in detection of Gram-negative foodborne pathogens. Microbiome 4 (1): 48. |
| ST66 | 3_2 | Pérez-Díaz, Ilenys M, Janet S Hayes, Eduardo Medina, Ashlee M Webber, Natasha Butz, Allison N Dickey, Zhongjing Lu, and Maria A Azcarate-Peril. 2019. Assessment of the Non-Lactic Acid Bacteria Microbiota in Fresh Cucumbers and Commercially Fermented Cucumber Pickles Brined with 6% NaCl. Food Microbiology 77 (February): 10–20. doi:10.1016/j.fm.2018.08.003. |
| ST67 | 3_2 | Delgado, S., Rachid, C. T. C. C., Fernández, E., Rychlik, T., Alegría, A,Peixoto, R. S., Mayo, B., 2013. Diversity of thermophilic bacteria in raw, pasteurized and selectively-cultured milk, as assessed by culturing, PCR-DGGE and pyrosequencing. Food Microbio. 36 (1): 103–111. |
| ST68 | 3_2 | Vepštaitė-Monstavičė, I., Lukša, J., Stanevičienė, R., Strazdaitė-Žielienė, Z., Yurchenko, V., Serva, S., Servienė, E., 2018. Distribution of apple and blackcurrant microbiota in Lithuania and the Czech Republic. Microb. Res. 206 (January): 1–8. |
| ST69 | 3_2 | Cruciata, M., Sannino, C., Ercolini, D., Scatassa, M. L., De Filippis, F., Mancuso, I., La Storia, A., Moschetti, G., Settanni, L., 2014. Animal rennets as sources of dairy lactic acid bacteria. Appl. Environ. Microbiol. 80 (7):2050–2061. |
| ST70 | 3_2 | Dees, M. W., Lysøe, E., Nordskog, B., Brurberg, M. B., 2015. Bacterial communities associated with surfaces of leafy greens: shift in composition and decrease in richness over time. Appl. Environ. Microbiol. 81 (4): 1530–1539. |
| ST71 | 3_2 | Jackson, C. R., Randolph, K. C., Osborn, S. L., Tyler, H. L., 2013. Culture dependent and independent analysis of bacterial communities associated with commercial salad leaf vegetables. BMC Microbiol. 13 (1): 274. |
| ST72 | 3_2 | Zhang, J. Li, Y., Liu, X., Lei, Y., Regenstein, J. M., Luo, Y. 2019. Characterization of the microbial composition and quality of lightly salted grass carp (Ctenopharyngodon Idellus) fillets with vacuum or Modified Atmosphere Packaging. Int. J. Food Microbiol. 293 (March) 87–93. |
| ST73 | 3_2 | Park, Eun-Jin, Jongsik Chun, Chang-Jun Cha, Wan-Soo Park, Che Ok Jeon, and Jin-Woo Bae. 2012. Bacterial Community Analysis During Fermentation of Ten Representative Kinds of Kimchi with Barcoded Pyrosequencing. Food Microbiology 30 (1): 197-204. doi:10.1016/j.fm.2011.10.011. |
| ST74 | 3_2 | Falardeau, J., Keeney, K., Trmčić, A., Kitts,D., Wang, S., 2019. Farm-to-fork profiling of bacterial communities associated with an artisan cheese production facility. Food Microbiol. 83: 48–58. |
| ST75 | 3_2 | Wang, H., Qin, X., Mi, S., Li, X., Wang, X., Yan, W., Zhang, C., 2019. Contamination of yellow-feathered broiler carcasses: microbial diversity and succession during processing. Food Microbiol. 83 (October): 18-26. |
| ST76 | 3_2 | Dharmarha, V., Pulido, N., Boyer,R. R., Pruden, A., Strawn, L. K., Ponder, M. A., 2018. Effect of post-harvest interventions on surficial carrot bacterial community dynamics, pathogen survival, and antibiotic resistance. Int. J. Food Microbiol. 291 (November): 25–34. |
| ST77 | 3_2 | Jarvis, K. G., White, J. R., Grim, C. J., Ewing, L., Ottesen, A. R., Beaubrun,J. J.-G., Pettengill, J. B., Brown,E., Hanes, D. E., 2015. Cilantro microbiome before and after nonselective pre-enrichment for Salmonella using 16S rRNA and metagenomic sequencing. BMC Microbiol. 15 (1): 160. |
| ST78 | 3_2 | Tatsika, S., Karamanoli, K., Karayanni, H., Genitsaris, S., 2019. Metagenomic characterization of bacterial communities on Ready-to-Eat vegetables and effects of household washing on their diversity and composition. Pathogens 8 (1): 37. |
| ST79 | 3_2 | Mota-Gutierrez, J. Botta, C., Ferrocino, I., Giordano, M., Bertolino, M., Dolci, P., Cannoni, M., Cocolin, L., 2018. Dynamics and biodiversity of bacterial and yeast communities during fermentation of cocoa beans. Appl. Environ. Microbiol. 84: 108–17. |
| ST80 | 3_2 | Ramezani, M., Hosseini, S. M., Ferrocino, I., Amoozegar, M. A., Cocolin, L., 2017. Molecular investigation of bacterial communities during the manufacturing and ripening of semi-hard Iranian Liqvan Cheese. Food Microbiol. 66: 64–71. |
| ST81 | 3_2 | Li, N., Wang, Y., You, C., Ren, J., Chen, W., Zheng, H., Liu, Z., 2018. Variation in raw milk microbiota throughout 12 months and the impact of weather conditions. Sci. Rep. 8: 2371. |
| ST82 | 3_2 | Angelopoulou, A., Holohan, R., Rea, M- C., Warda, A. K., Hill, C., Ross, R. P., 2019. Bovine mastitis Is a polymicrobial disease requiring a polydiagnostic approach. int. Dairy J. 99: 104539. |
| ST83 | 3_2 | Kim, H.-E., Lee, J.-J., Lee, M.-J., Kim, B.-S., 2019. Analysis of microbiome in raw chicken meat from butcher shops and packaged products in South Korea to detect the potential risk of foodborne illness. Frin 122: 517–527. |
| ST84 | 3_2 | Osimani, A., Ferrocino, I., Agnolucci,M., Cocolin, L., Giovannetti, M., Cristani, C., Palla, M., Milanović, V., Roncolini, A., Sabbatini, R., Garofalo, C., Clementi, F., Cardinali, F., Petruzzelli, A., Gabucci, C., Tonucci, F., Aquilanti, L., 2019. Unveiling Hákarl: a study of the microbiota of the traditional Icelandic fermented fish. Food Microbiol. 82: 560–572. |
| ST85 | 3_2 | Tan, X., Chung, T., Chen, Y., Macarisin, D., LaBorde, L., Kova, J., 2019. The occurrence of Listeria monocytogenes Is associated with built environment microbiota in three tree fruit processing facilities. Microbiome 7 (1): 115–118. |
| ST86 | 3_2 | Oshiro, M., Momoda, R., Tanaka, M., Zendo, T., Nakayama, J., 2019. Dense tracking of the dynamics of the microbial community and chemicals constituents in spontaneous wheat sourdough during two months of backslopping. J. Biosci. Bioeng. 128: 170–176. |
| ST87 | 3_2 | Kable, M. E., Srisengfa, Y., Xue, Z., Coates, L. C., Marco, M. L., 2019. Viable and total bacterial populations undergo equipment- and time-dependent shifts during milk processing. Appl. Environ. Microbiol. 85: 190. |
| ST88 | 3_2 | Castro, I., Alba, C., Aparicio, M., Arroyo,R., Jiménez, L., Fernández, L., Arias, R., Rodriguez, J. M., 2019. Metataxonomic and immunological analysis of milk from ewes with or without a history of mastitis. J. Dairy Sci. 102: 9298–9311. |
| ST89 | 3_2 | Catozzi, C., Bonastre, A. S., Francino, O., Lecchi, C., De Carlo, E., Vecchio, D., Martucciello, A., Fraulo, P., Bronzo, V., Cuscó, A., D’Andreano, S., Ceciliani, F., 2017. The microbiota of water buffalo milk during mastitis. PLoS One 12 (9): e0184710. |
| ST90 | 3_2 | Antunes-Rohling, A., Calero, S., Halaihel, N., Marquina, P., Raso, J., Calanche,J., Beltrán, J. A., Álvarez, Cebrián, G., 2019. Characterization of the spoilage microbiota of Hake fillets packaged under a Modified Atmosphere (MAP) rich in CO2 (50% CO2/50% N2) and stored at different temperatures. Foods 8 (489): 1–14. |
| ST91 | 3_2 | Vandeweyer, D., Crauwels, S., Lievens,B., Van Campenhout, L., 2017. Metagenetic analysis of the bacterial communities of edible insects from diverse production cycles at industrial rearing companies. Int. J. Food Microbiol. 261: 11–18. |
| ST92 | 3_2 | Vandeweyer, D., Wynants, E., Crauwels, S., Verreth, C., Viaene,N., Claes, J., Lievens, B., Van Campenhout, L., 2018. Microbial dynamics during industrial rearing, processing, and storage of the tropical house cricket (Gryllodes sigillatus) for human consumption. Appl. Environ. Microbiol. 84 (12): e00255–18. |
| ST93 | 3_2 | Wynants, E., Crauwels, S., Verreth, C., Gianotten, N., Lievens, B., Claes, J., Van Campenhout, L., 2018. Microbial dynamics during production of lesser mealworms (Alphitobius diaperinus) for human consumption at industrial scale. Food Microbiol. 70 (April): 181–191. |
| ST94 | 3_2 | Keshri, J., Krouptiski, Y., Abu-Fani, L., Achmon, Y., Bauer, T. S., Zarka, O., Maler, I., Pinto, R., Sela Saldinger, S., 2019. Dynamics of bacterial communities in Alfalfa and Mung Bean sprouts during refrigerated conditions. Food Microbiol. 84: 103261. |
| ST95 | 3_2 | Diaz, M., Kellingray, L., Akinyemi, N., Olaoluwa Adefiranye, O., Olaonipekun, A. B., Bayili,G., R., Ibezim, J., du Plessis, A. S., Houngbédji, M., Kamya, D., Mukisa, I. M., Mulaw, G., Josiah, S. M., Chienjo, W. O., Atter, A., Agbemafle, E., Annan, T., Ackah, N. b:, Buys, E. M., Hounhouigan, D. J., Muyanja, C., Nakavuma, J., Odeny, D. A., Sawadogo-Lingani, H., Tefera, A. T., Amoa-Awua, W., Obodai, M., Mayer, M. J., Oguntoyinbo, F. A., Narbad, A., 2019. Comparison of the microbial composition of African fermented foods using amplicon sequencing. Sci. Rep. 9 (1): 1–8. |
| ST96 | 3_2 | Vahdatzadeh, M., Deveau, A., Splivallo, R., 2019. Are bacteria responsible for aroma deterioration upon storage of the Black Truffle Tuber aestivum: a microbiome and volatilome study. Food Microbiol. 84 (December): 103251 |
| ST97 | 3_2 | De Filippis, F., Aponte, M., Piombino, P., Lisanti, M. T., Moio, L., Ercolini, D., Blaiotta, G., 2019. Influence of microbial communities on the chemical and sensory features of Falanghina sweet passito wines. Food Res. Int. 120 (June): 740–47. |
| ST98 | 3_2 | Astudillo-Melgar, F., Ochoa-Leyva, A., Utrilla, J., Huerta-Beristain, G., 2019. Bacterial diversity and population dynamics during the fermentation of palm wine from Guerrero Mexico. Front. Microbiol. 10: 531. |
| ST99 | 3_2 | Du, F., Zhang, X., Gu,H., Song, J., Gao, X., 2019. Dynamic changes in the bacterial community during the fermentation of Traditional Chinese Fish Sauce (TCFS) and their correlation with TCFS quality. Microorganisms 7 (9): 371. |
| ST100 | 3_2 | Jampaphaeng, Krittanon, Ilario Ferrocino, Manuela Giordano, Kalliopi Rantsiou, Suppasil Maneerat, and Luca Cocolin. 2018. Microbiota Dynamics and Volatilome Profile During Stink Bean Fermentation (Sataw-Dong) with Lactobacillus plantarum KJ03 as a Starter Culture. Food Microbiology 76 (December): 91–102. doi:10.1016/j.fm.2018.04.012. |
| ST101 | 3_2 | Li, N., Zhang, Y., Wu, Q., Gu, Q., Chen, M., Zhang, Y., Sun, X., Zhang, J., 2019. High-throughput sequencing analysis of bacterial community composition and quality characteristics in refrigerated pork during storage. Food Microbiol. 83 (October): 86–94. |
| ST102 | 3_2 | Lv, J., Yang, Z., Xu, W., Li, S., Liang, H., Ji, C., Yu, C., Zhu, B., Lin, X., 2019. Relationships between bacterial community and metabolites of sour meat at different temperature during the fermentation. Int. J. Food Microbiol. 307 (October): 108286. |
| ST103 | 3_2 | Wang, J., Wang, R., Xiao, Q., Liu, C., Jiang, L., Deng, F., Zhou, H., 2019. Analysis of bacterial diversity during fermentation of Chinese traditional fermented chopped pepper. Lett. Appl. Microbiol. 69 (5): 346–352. |
| ST104 | 3_2 | Hauptmann, A. L., Paulov, P., Castro-Mej’a, J. L., Hansen, L- H., Sicheritz-Pontén, T., Mulvad, G., Nielsen, D. S., 2020. The microbial composition of dried fish prepared according to Greenlandic Inuit traditions and industrial counterparts. Food Microbiol. 85 (February): 103305. |
| ST105 | 3_2 | Motato, K., Milani, C., Ventura, M., Valencia, F., Ruas-Madiedo, P., Delgado, S., 2017. Bacterial diversity of the Colombian fermented milk “Suero Costeño” assessed by culturing and high-throughput sequencing and DGGE analysis of 16S rRNA gene amplicons. Food Microbiol. 68: 129-136. https://dx.doi.org/10.1016/j.fm.2017.07.011 |
| ST106 | 3_2 | De Pasquale, I., Cagno, R., Buchin, S., De Angelis, M., Gobbetti, M., 2016. Spatial distribution of the metabolically active microbiota within Italian PDO ewes' milk cheeses. PLoS One 11(4): e0153213. https://dx.doi.org/10.1371/journal.pone.0153213 |
| ST107 | 3_2 | Kamimura, B., Cabral, L., Noronha, M., Baptista, R., Nascimento, H., Sant'Ana, A., 2020. Amplicon sequencing reveals the bacterial diversity in milk, dairy premises and Serra da Canastra artisanal cheeses produced by three different farms. Food Microbiol. 89(): 103453. https://dx.doi.org/10.1016/j.fm.2020.103453 |
| ST108 | 3_2 | Levante, A., Filippis, F., Storia, A., Gatti, M., Neviani, E., Ercolini, D., Lazzi, C.,(2017. Metabolic gene-targeted monitoring of non-starter lactic acid bacteria during cheese ripening. Int. Journal Food Microbiol. 257: 276-284. https://dx.doi.org/10.1016/j.ijfoodmicro.2017.07.002 |
| ST109 | 3_2 | Parker, M., Zobrist, S., Donahue, C., Edick, C., Mansen, K., Nadjari, M., Heerikhuisen, M., Sybesma, W., Molenaar, D., Diallo, A., Milani, P., Kort, R., 201). Naturally fermented milk from Northern Senegal: bacterial community composition and probiotic enrichment with Lactobacillus rhamnosus. Front. Microbiol. 9(): 2218. https://dx.doi.org/10.3389/fmicb.2018.02218 |
| ST110 | 3_2 | Calasso, M., Minervini, F., Filippis, F., Ercolini, D., Angelis, M., Gobbetti, M., 2020. Attenuated Lactococcus lactis and surface bacteria as tools for conditioning the microbiota and driving the ripening of semisoft Caciotta cheese. Appl. Environ. Microbiol. 86(5):e02165-19 https://dx.doi.org/10.1128/aem.02165-19 |
| ST111 | 3_2 | Carafa, I., Stocco, G., Nardin, T., Larcher, R., Bittante, G., Tuohy, K., Franciosi, E., 2019. Production of naturally g-aminobutyric acid-enriched cheese using the dairy strains Streptococcus thermophilus 84C and Lactobacillus brevis DSM 32386. Front. Microbiol. 10:9 doi:10.3389/fmicb.2019.00093. |
| ST112 | 3_2 | Wang, H., Qin, X., Li, X., Wang, X., Gao, H., Zhang, C., 2019. Changes in the microbial communities of air- and water-chilled yellow-feathered broilers during storage at 2 °C Food Microbiol. 87:103390 https://dx.doi.org/10.1016/j.fm.2019.103390 |
| ST113 | 3_2 | Pini, F., Aquilani, C., Giovannetti, L., Viti, C., Pugliese, C., 2020. Characterization of the microbial community composition in Italian Cinta Senese sausages dry-fermented with natural extracts as alternatives to sodium nitrite. Food Microbiol. 89(), 103417. https://dx.doi.org/10.1016/j.fm.2020.103417 |
| ST114 | 3_2 | Settanni, L., Barbaccia, P., Bonanno, A., Ponte, M., Gerlando, R., Franciosi, E., Grigoli, A., Gaglio, R., 2020. Evolution of indigenous starter microorganisms and physicochemical parameters in spontaneously fermented beef, horse, wild boar and pork salamis produced under controlled conditions. Food Microbiol. 87(): 103385. https://dx.doi.org/10.1016/j.fm.2019.103385 |
| ST115 | 3_2 | Choi, J., Lee, S., Rackerby, B., Goddik, L., Frojen, R., Ha, S., Kim, J., Park, S., 2020. Microbial communities of a variety of cheeses and comparison between core and rind region of cheeses. J. Dairy Sci. 103: in press https://dx.doi.org/10.3168/jds.2019-17455 |
| ST116 | 3_2 | Mu, Y., Su, W., Mu, Y., Jiang, L. (2019). Combined Application of High-Throughput Sequencing and Metabolomics Reveals Metabolically Active Microorganisms During Panxian Ham Processing. Frontiers in Microbiology 10(), 3012. https://dx.doi.org/10.3389/fmicb.2019.03012 |
| ST117 | 3_2 | Pérez-Díaz, I., Dickey, A., Fitria, R., Ravishankar, N., Hayes, J., Campbell, K., Arritt, F. (2020). Modulation of the bacterial population in commercial cucumber fermentations by brining salt type. J. Appl. Microbiol., https://dx.doi.org/10.1111/jam.14597 |
| ST118 | 3_2 | Boreczek, J., Litwinek, D., Żylińska-Urban, J., Izak, D., Buksa, K., Gawor, J., Gromadka, R., Bardowski, J., Kowalczyk, M., 2020. Bacterial community dynamics in spontaneous sourdoughs made from wheat, spelt, and rye wholemeal flour. MicrobiologyOpen 2020;00:e1009 https://dx.doi.org/10.1002/mbo3.1009 |
| ST119 | 3_2 | Wassermann, B., Kusstatscher, P., Berg, G.,2019. Microbiome response to hot water treatment and potential synergy with biological control on stored apples. Frontiers in Microbiology 10(), 2502. https://dx.doi.org/10.3389/fmicb.2019.02502 |
| ST120 | 3_2 | Yang, H., Yang, L., Zhang, J., Li, H., Tu, Z., Wang, X., 2019. Exploring functional core bacteria in fermentation of a traditional Chinese food, Aspergillus-type douchi. PloS one 14(12), e0226965. |
| ST121 | 3_2 | Belleggia, L., Aquilanti, L., Ferrocino, I., Milanović, V., Garofalo, C., Clementi, F., Cocolin, L., Mozzon, M., Foligni, R., Haouet, M., Scuota, S., Framboas, M., Osimani, A., 2020. Discovering microbiota and volatile compounds of surströmming, the traditional Swedish sour herring. Food Microbiol. 91(), 103503. https://dx.doi.org/10.1016/j.fm.2020.103503 |
| ST122 | 3_2 | Sørensen, J., Bøknæs, N., Mejlholm, O., Dalgaard, P., 2020. Superchilling in combination with modified atmosphere packaging resulted in long shelf-life and limited microbial growth in Atlantic cod (Gadus morhua L.) from capture-based-aquaculture in Greenland. Food Microbiol. 88(), 103405. https://dx.doi.org/10.1016/j.fm.2019.103405 |
| ST123 | 3_2 | Catozzi, C., Ceciliani, F., Lecchi, C., Talenti, A., Vecchio, D., Carlo, E., Grassi, C., Sánchez, A., Francino, O., Cuscó, A., 2020. Short communication: Milk microbiota profiling on water buffalo with full-length 16S rRNA using nanopore sequencing. J. Dairy Sci. 103(3), 2693-2700. https://dx.doi.org/10.3168/jds.2019-17359 |
| ST124 | 3_2 | Catozzi, C., Ceciliani, F., Lecchi, C., Talenti, A., Vecchio, D., Carlo, E., Grassi, C., Sánchez, A., Francino, O., Cuscó, A., 2020. Short communication: Milk microbiota profiling on water buffalo with full-length 16S rRNA using nanopore sequencing. J. Dairy Sci. 103(3), 2693-2700. https://dx.doi.org/10.3168/jds.2019-17359 |
| ST125 | 3_2 | McHugh, A., Feehily, C., Fenelon, M., Gleeson, D., Hill, C., Cotter, P., 2020. Tracking the dairy microbiota from farm bulk tank to skimmed milk powder. mSystems: 5(2) e00262-20 https://dx.doi.org/10.1128/msystems.00226-20 |
| ST126 | 3_2 | Maoloni, A., Blaiotta, G., Ferrocino, I., Mangia, N., Osimani, A., Milanović, V., Cardinali, F., Cesaro, C., Garofalo, C., Clementi, F., Pasquini, M., Trombetta, M., Cocolin, L., Aquilanti, L., 2020. Microbiological characterization of Gioddu, an Italian fermented milk. Int. J. Food Microbiol. 323(): 108610. https://dx.doi.org/10.1016/j.ijfoodmicro.2020.108610 |
| ST127 | 3_2 | Gaglio, R., Cirlincione, F., Miceli, G., Franciosi, E., Gerlando, R., Francesca, N., Settanni, L., Moschetti, G., 2020. Microbial dynamics in durum wheat kernels during aging. Int. J. Food Microbiol 324(): 108631. https://dx.doi.org/10.1016/j.ijfoodmicro.2020.108631 |
| ST128 | 3_2 | Zagdoun, M., Coeuret, G., N'Dione, M., Champomier-Vergès, M., Chaillou, S., 2020. Large microbiota survey reveals how the microbial ecology of cooked ham is shaped by different processing steps. Food Microbiol. 91(): 103547. https://dx.doi.org/10.1016/j.fm.2020.103547 |
| ST129 | 3_2 | Tan, G., Hu, M., Li, X., Pan, Z., Li, M., Li, L., Yang, M., 2020. High-Throughput Sequencing and metabolomics reveal differences in bacterial diversity and metabolites between Red and White Sufu. Front. Microbiol. 11(): 758. https://dx.doi.org/10.3389/fmicb.2020.00758 |
| ST130 | 3_2 | Choi, H., Hwang, B., Kim, B., Choi, S., 2020. Influence of pathogen contamination on beef microbiota under different storage temperatures. Food Res. Int. 132(): 109118. https://dx.doi.org/10.1016/j.foodres.2020.109118 |
| ST131 | 3_2 | Choi, J., Lee, S., Rackerby, B., Frojen, R., Goddik, L., Ha, S., Park, S., 2020. Assessment of overall microbial community shift during Cheddar cheese production from raw milk to aging. Appl. Microbiol. Biotechnol. 104(14) 6249-6260 https://dx.doi.org/10.1007/s00253-020-10651-7 |
| ST132 | 3_2 | Nalepa, B., Ciesielski, S., Aljewicz, M., 2020. The microbiota of Edam cheeses determined by cultivation and High-Throughput Sequencing of the 16S rRNA amplicon. Appl. Sci. 10(12): 4063. https://dx.doi.org/10.3390/app10124063 |
| ST133 | 3_2 | Anagnostopoulos, D., Kamilari, E., Tsaltas, D., 2020. Evolution of bacterial communities, physicochemical changes and sensorial attributes of natural whole and cracked Picual table olives during spontaneous and inoculated fermentation. Front. microbiol. 11(), 1128. https://dx.doi.org/10.3389/fmicb.2020.01128 |
| ST134 | 3_2 | Carafa, I., Navarro, I., Bittante, G., Tagliapietra, F., Gallo, L., Tuohy, K., Franciosi, E., 2020. Shift in the cow milk microbiota during alpine pasture as analyzed by culture dependent and high-throughput sequencing techniques Food Microbiology 91(), 103504. https://dx.doi.org/10.1016/j.fm.2020.103504 |
| ST135 | 3_2 | Luzzi, G., Brinks, E., Fritsche, J., Franz, C., 2020. Microbial composition of sweetness-enhanced yoghurt during fermentation and storage. AMB Express 10(1): 131. https://dx.doi.org/10.1186/s13568-020-01069-5 |
| ST136 | 3_2 | Dolci, P., Ferrocino, I., Giordano, M., Pramotton, R., Vernetti-Prot, L., Zenato, S., Barmaz, A., 2020. Impact of Lactococcus lactis as starter culture on microbiota and metabolome profile of an Italian raw milk cheese. Int. Dairy J. 110(): 104804. https://dx.doi.org/10.1016/j.idairyj.2020.104804 |
| ST137 | 3_2 | Comasio, A., Verce, M., Kerrebroeck, S., Vuyst, L., 2020. Diverse microbial composition of sourdoughs from different origin.s Front. Microbiol. 11(): 1212. https://dx.doi.org/10.3389/fmicb.2020.01212 |
| ST138 | 3_2 | Hwang, B., Choi, H., Choi, S., Kim, B, 2020. Analysis of microbiota structure and potential functions influencing spoilage of fresh beef meat. Front. Microbiol. 11(): 1657. https://dx.doi.org/10.3389/fmicb.2020.01657 |
| ST139 | 3_2 | Kang, S., Ravensdale, J., Coorey, R., Dykes, G., Barlow, R., 2020. Bacterial community analysis using 16S rRNA amplicon sequencing in the boning room of Australian beef export abattoirs. Int. J. Food Microbiol. 332(): 108779. https://dx.doi.org/10.1016/j.ijfoodmicro.2020.108779 |
| ST140 | 3_2 | Kang, S., Ravensdale, J., Coorey, R., Dykes, G. A. & Barlow, R., 2019. A comparison of 16S rRNA profiles through slaughter in Australian export beef abattoirs. Front. Microbiol. 10: 2747. https://doi.org/10.3389/fmicb.2019.00093 |
| ST141 | 3_2 | Penland, M., Deutsch, S., Falentin, H., Pawtowski, A., Poirier, E., Visenti, G., Meur, C., Maillard, M., Thierry, A., Mounier, J., Coton, M., 2020. Deciphering microbial community dynamics and biochemical changes during Nyons black olive natural fermentations. Front. Microbiol. 11(): 586614. https://doi.org/10.3389/fmicb.2019.00093 |
| ST142 | 3_2 | Chen, X., Zhu, L., Liang, R., Mao, Y., Hopkins, D., Li, K., Dong, P., Yang, X., Niu, L., Zhang, Y., Luo, X., 2019. Shelf-life and bacterial community dynamics of vacuum packaged beef during long-term super-chilled storage sourced from two Chinese abattoirs. Food Res. Int. 130(): 108937. https://doi.org/10.3389/fmicb.2019.00093 |
| ST143 | 3_2 | Han, D., Chun, B., Feng, T., Kim, H., Jeon, C., 2020. Dynamics of microbial communities and metabolites in ganjang, a traditional Korean fermented soy sauce, during fermentation. Food Microbiol. 92(): 103591. https://doi.org/10.3389/fmicb.2019.00093 |
| ST144 | 3_2 | Lin, L., Wu, J., Chen, X., Huang, L., Zhang, X., Gao, X., 2020. The role of the bacterial community in producing a peculiar smell in Chinese fermented sour soup. Microorganisms, 8(9): 1270. https://dx.doi.org/10.3390/microorganisms8091270 |
| ST145 | 3_2 | Zheng, X., Ge, Z., Lin, K., Zhang, D., Chen, Y., Xiao, J., Wang, B., Shi, X. (2020). Dynamic changes in bacterial microbiota succession and flavour development during milk fermentation of Kazak artisanal cheese International Dairy Journal 113(), 104878. https://dx.doi.org/10.1016/j.idairyj.2020.104878 |
| ST146 | 3_2 | Castellanos-Rozo, J., Pulido, R., Grande, M. J., Lucas, R., Gálvez, A., 2020. Analysis of the bacterial diversity of Paipa cheese (a traditional raw cow's milk cheese from Colombia) by High-Throughput Sequencing. Microorganisms, 8(2): 218. https://dx.doi.org/10.3390/microorganisms8020218 |
| ST147 | 3_2 | Hou, K., Tong, J., Zhang, H., Gao, S., Guo, Y., Niu, H., Xiong, B., Jiang, L., 2020. Microbiome and metabolic changes in milk in response to artemisinin supplementation in dairy cows. AMB Expr., 10(1): 154. https://dx.doi.org/10.1186/s13568-020-01080-w |
| ST148 | 3_2 | Bottari, B., Levante, A., Bancalari, E., Sforza, S., Bottesini, C., Prandi, B., Filippis, F., Ercolini, D., Nocetti, M., Gatti, M., 2020. The Interrelationship between microbiota and peptides during ripening as a driver for Parmigiano Reggiano cheese quality. Front. Microbiol. 11(): 581658. https://dx.doi.org/10.3389/fmicb.2020.581658 |
| ST149 | 3_2 | Cremonesi, P., Morandi, S., Ceccarani, C., Battelli, G., Castiglioni, B., Cologna, N., Goss, A., Severgnini, M., Mazzucchi, M., Partel, E., Tamburini, A., Zanini, L., Brasca, M., 2020. Raw milk microbiota modifications as affected by chlorine usage for cleaning procedures: the Trentingrana PDO case. Front. Microbiol. 11(): 564749. https://dx.doi.org/10.3389/fmicb.2020.564749 |
| ST150 | 3_2 | Biolcati, F., Ferrocino, I., Bottero, M. T. & Dalmasso, A., 2020. Short communication: High-throughput sequencing approach to investigate Italian artisanal cheese production. J Dairy Sci 103: 10015–10021. |
| ST151 | 3_2 | Doster, E., Thomas, K.M., Weinroth, M.D., Parker, J.K., Crone, K.K., Arthur, T.M., Schmidt, J.W., Wheeler, T.L., Belk, K.E., Morley, P.S., 2020. Metagenomic characterization of the microbiome and resistome of retail ground beef products. Front Microbiol 11: 541972. |
| ST152 | 3_2 | Zwirzitz, B., Wetzels, S.U., Dixon, E.D., Stessl, B., Zaiser, A., Rabanser, I., Thalguter, S., Pinior, B., Roch, F.-F., Strachan, C., Zanghellini, J., Dzieciol, M., Wagner, M., Selberherr, E., 2020. The sources and transmission routes of microbial populations throughout a meat processing facility. Npj Biofilms Microbiomes: 6, 26. |
| ST153 | 3_2 | Kaur, M., Williams, M., Bissett, A., Ross, T., Bowman, J.P., 2021. Effect of abattoir, livestock species and storage temperature on bacterial community dynamics and sensory properties of vacuum packaged red meat. Food Microbiol: 94, 103648. |
| ST154 | 3_2 | Ding, R., Liu, Yiming, Yang, S., Liu, Yumeng, Shi, H., Yue, X., Wu, R., Wu, J., 2020. High-throughput sequencing provides new insights into the roles and implications of core microbiota present in pasteurized milk. Food Res Int 137: 109586. https://doi.org/10.1016/j.foodres.2020.109586 |
| ST155 | 3_2 | Luoyizha, W., Wu, X., Zhang, M., Guo, X., Li, H., Liao, X., 2020. Compared analysis of microbial diversity in donkey milk from Xinjiang and Shandong of China through High-throughput sequencing. Food Res Int 137: 109684. https://doi.org/10.1016/j.foodres.2020.109684 |
| ST156 | 3_2 | Frigerio, J., Agostinetto, G., Galimberti, A., Mattia, F.D., Labra, M., Bruno, A., 2020. Tasting the differences: Microbiota analysis of different insect-based novel food. Food Res Int 137: 109426. https://doi.org/10.1016/j.foodres.2020.109426 |
| ST157 | 3_2 | Afshari, R., Pillidge, C.J., Dias, D.A., Osborn, A.M., Gill, H., 2020. Microbiota and Metabolite Profiling Combined With Integrative Analysis for Differentiating Cheeses of Varying Ripening Ages. Front Microbiol 11, 592060. https://doi.org/10.3389/fmicb.2020.592060 |
| ST158 | 3_2 | Garofalo, C., Ferrocino, I., Reale, A., Sabbatini, R., Milanović, V., Alkić-Subašić, M., Boscaino, F., Aquilanti, L., Pasquini, M., Trombetta, M.F., Tavoletti, S., Coppola, R., Cocolin, L., Blesić, M., Sarić, Z., Clementi, F., Osimani, A., 2020. Study of kefir drinks produced by backslopping method using kefir grains from Bosnia and Herzegovina: Microbial dynamics and volatilome profile. Food Res Int 137: 109369. https://doi.org/10.1016/j.foodres.2020.109369 |
| ST159 | 3_2 | Chen, X., Zhao, J., Zhu, L., Luo, X., Mao, Y., Hopkins, D.L., Zhang, Y., Dong, P., 2020. Effect of modified atmosphere packaging on shelf life and bacterial community of roast duck meat. Food Res Int 137: 109645. https://doi.org/10.1016/j.foodres.2020.109645 |
| ST160 | 3_2 | Østlie, H.M., Porcellato, D., Kvam, G., Wicklund, T., 2021. Investigation of the microbiota associated with ungerminated and germinated Norwegian barley cultivars with focus on lactic acid bacteria. Int J Food Microbiol 341: 109059. https://doi.org/10.1016/j.ijfoodmicro.2021.109059 |
| ST161 | 3_2 | Tyakht, A., Kopeliovich, A., Klimenko, N., Efimova, D., Dovidchenko, N., Odintsova, V., Kleimenov, M., Toshchakov, S., Popova, A., Khomyakova, M., Merkel, A., 2021. Characteristics of bacterial and yeast microbiomes in spontaneous and mixed-fermentation beer and cider. Food Microbiol 94: 103658. https://doi.org/10.1016/j.fm.2020.103658 |
| ST162 | 3_2 | Rocha-Arriaga, C., Espinal-Centeno, A., Martinez-Sánchez, S., Caballero-Pérez, J., Alcaraz, L.D., Cruz-Ramírez, A., 2020. Deep microbial community profiling along the fermentation process of pulque, a biocultural resource of Mexico. Microbiol Res 241, 126593. https://doi.org/10.1016/j.micres.2020.126593 |
| ST163 | 3_2 | Rault, L., Lévêque, P.-A., Barbey, S., Launay, F., Larroque, H., Loir, Y.L., Germon, P., Guinard-Flament, J., Even, S., 2020. Bovine teat cistern microbiota composition and richness are associated with the immune and microbial responses during transition to once-daily milking. Front Microbiol 11: 602404. https://doi.org/10.3389/fmicb.2020.602404 |
| ST164 | 3_2 | Dean, C.J., Slizovskiy, I.B., Crone, K.K., Pfennig, A.X., Heins, B.J., Caixeta, L.S., Noyes, N.R., 2021. Investigating the cow skin and teat canal microbiomes of the bovine udder using different sampling and sequencing approaches. J Dairy Sci 104: 644–661. https://doi.org/10.3168/jds.2020-18277 |
| ST165 | 3_2 | Penland, M., Falentin, H., Parayre, S., Pawtowski, A., Maillard, M.-B., Thierry, A., Mounier, J., Coton, M., Deutsch, S.-M., 2021. Linking Pélardon artisanal goat cheese microbial communities to aroma compounds during cheese-making and ripening. Int J Food Microbiol 345: 109130. https://doi.org/10.1016/j.ijfoodmicro.2021.109130 |
| ST166 | 3_2 | Bossaert, S., Winne, V., Opstaele, F.V., Buyse, J., Verreth, C., Herrera-Malaver, B., Geel, M.V., Verstrepen, K.J., Crauwels, S., Rouck, G.D., Lievens, B., 2021. Description of the temporal dynamics in microbial community composition and beer chemistry in sour beer production via barrel ageing of finished beers. Int J Food Microbiol 339, 109030. https://doi.org/10.1016/j.ijfoodmicro.2020.109030 |
| ST167 | 3_2 | Wen, R., Lv, Y., Li, X., Chen, Q., Kong, B., 2021. High-throughput sequencing approach to reveal the bacterial diversity of traditional yak jerky from the Tibetan regions. Meat Sci 172, 108348. https://doi.org/10.1016/j.meatsci.2020.108348 |
| ST168 | 3_2 | Botta, C., Ferrocino, I., Pessione, A., Cocolin, L., Rantsiou, K., 2020. Spatiotemporal Distribution of the Environmental Microbiota in Food Processing Plants as Impacted by Cleaning and Sanitizing Procedures: the Case of Slaughterhouses and Gaseous Ozone. Appl Environ Microb 86. https://doi.org/10.1128/aem.01861-20 |
| ST169 | 3_2 | Porcellato, D., Meisal, R., Bombelli, A., Narvhus, J.A., 2020. A core microbiota dominates a rich microbial diversity in the bovine udder and may indicate presence of dysbiosis. Sci Rep-uk 10, 21608. https://doi.org/10.1038/s41598-020-77054-6 |
| ST170 | 3_2 | Porcellato, D., Smistad, M., Bombelli, A., Abdelghani, A., Jørgensen, H.J., Skeie, S.B., 2021. Longitudinal Study of the Bulk Tank Milk Microbiota Reveals Major Temporal Shifts in Composition. Front Microbiol 12, 616429. https://doi.org/10.3389/fmicb.2021.616429 |
| ST171 | 4_1 | Zhadyra, S., Han, X., Anapiyayev, B.B., Tao, F., Xu, P., 2021. Bacterial diversity analysis in Kazakh fermented milks Shubat and Ayran by combining culture-dependent and culture-independent methods. Lwt 141, 110877. https://doi.org/10.1016/j.lwt.2021.110877 |
| ST172 | 4_1 | Salazar, J.K., Gonsalves, L.J., Fay, M., Ramachandran, P., Schill, K.M., Tortorello, M.L., 2021. Metataxonomic Profiling of Native and Starter Microbiota During Ripening of Gouda Cheese Made With Listeria monocytogenes-Contaminated Unpasteurized Milk. Front Microbiol 12, 642789. https://doi.org/10.3389/fmicb.2021.642789 |
| ST173 | 4_1 | Sawada, K., Koyano, H., Yamamoto, N., Yamada, T., 2021. The relationships between microbiota and the amino acids and organic acids in commercial vegetable pickle fermented in rice-bran beds. Sci Rep-uk 11, 1791. https://doi.org/10.1038/s41598-021-81105-x |
| ST174 | 4_1 | Unno, R., Suzuki, T., Matsutani, M., Ishikawa, M., 2021. Evaluation of the Relationships Between Microbiota and Metabolites in Soft-Type Ripened Cheese Using an Integrated Omics Approach. Front Microbiol 12, 681185. https://doi.org/10.3389/fmicb.2021.681185 |
| ST175 | 4_1 | Ryu, S., Park, W.S., Yun, B., Shin, M., Go, G., Kim, J.N., Oh, S., Kim, Y., 2021. Diversity and characteristics of raw milk microbiota from Korean dairy farms using metagenomic and culturomic analysis. Food Control 127, 108160. https://doi.org/10.1016/j.foodcont.2021.108160 |
| ST176 | 4_1 | Mancini, A., Rodriguez, M.C., Zago, M., Cologna, N., Goss, A., Carafa, I., Tuohy, K., Merz, A., Franciosi, E., 2021. Massive Survey on Bacterial–Bacteriophages Biodiversity and Quality of Natural Whey Starter Cultures in Trentingrana Cheese Production. Front Microbiol 12, 678012. https://doi.org/10.3389/fmicb.2021.678012 |
| ST177 | 4_1 | Irlinger, F., Monnet, C., 2021. Temporal differences in microbial composition of Époisses cheese rinds during ripening and storage. J Dairy Sci 104, 7500–7508. https://doi.org/10.3168/jds.2021-20123 |
| ST178 | 4_1 | Nikoloudaki, O., Junior, W.J.F.L., Borruso, L., Campanaro, S., Angelis, M.D., Vogel, R.F., Cagno, R.D., Gobbetti, M., 2021. How multiple farming conditions correlate with the composition of the raw cow’s milk lactic microbiome. Environ Microbiol. https://doi.org/10.1111/1462-2920.15407 |
| ST179 | 4_1 | Wilson, A., Chandry, P.S., Turner, M.S., Courtice, J.M., Fegan, N., 2021. Comparison between cage and free-range egg production on microbial composition, diversity and the presence of Salmonella enterica. Food Microbiol 97, 103754. https://doi.org/10.1016/j.fm.2021.103754 |
| ST180 | 4_1 | Yu, M., Wang, X., Yan, A., 2021. Microbial Profiles of Retail Pacific Oysters (Crassostrea gigas) From Guangdong Province, China. Front Microbiol 12, 689520. https://doi.org/10.3389/fmicb.2021.689520 |

**Supplementary Table 4**. Median and 90° percentile values for the frequency of taxonomic assignment of ASVs at the genus level or below in FoodMicrobionet studies from 34 to 180, by 16S gene target region (FALSE or TRUE indicate if the sequences showed overlapping; if not species level assignment was not performed). n indicates the number of studies using a given target. Both the absolute frequency and the frequency weighted by sequence abundance are shown.

|  |  | Frequency of genus + species identification | | Frequency of genus + species identification, weighted by sequence abundance | |
| --- | --- | --- | --- | --- | --- |
| Target | n | median | 90° perc | median | 90° perc |
| V1-V2_TRUE | 3 | 0.804 | 0.878 | 0.859 | 0.910 |
| V1-V3_FALSE | 5 | 0.767 | 0.802 | 0.969 | 0.985 |
| V1-V3_TRUE | 14 | 0.863 | 0.965 | 0.992 | 0.999 |
| V3_TRUE | 3 | 0.734 | 0.841 | 0.839 | 0.934 |
| V3-V4_FALSE | 28 | 0.745 | 0.803 | 0.945 | 0.994 |
| V3-V4_TRUE | 64 | 0.857 | 0.944 | 0.980 | 0.999 |
| V4_TRUE | 25 | 0.795 | 0.867 | 0.887 | 0.991 |
| V4-V5_FALSE | 1 | 0.771 | 0.771 | 0.305 | 0.305 |
| V4-V5_TRUE | 1 | 0.898 | 0.898 | 0.938 | 0.938 |
| V5-V6_TRUE | 1 | 0.787 | 0.787 | 0.866 | 0.866 |
| V5-V9_TRUE | 1 | 0.640 | 0.640 | 0.923 | 0.923 |
| V6-V8_TRUE | 1 | 0.871 | 0.871 | 0.964 | 0.964 |

**Supplementary figure 3.** Box and jitter plots showing the unweighted (sequence relative abundance ignored) distribution of frequencies of taxonomic assignments at the genus level or below in FoodMicrobionet studies 34 to 180. The average values for the number of issues encountered during bioinformatic processing (high sequence losses during filtering or chimera removal, low number of final sequences, low diversity) is also shown.

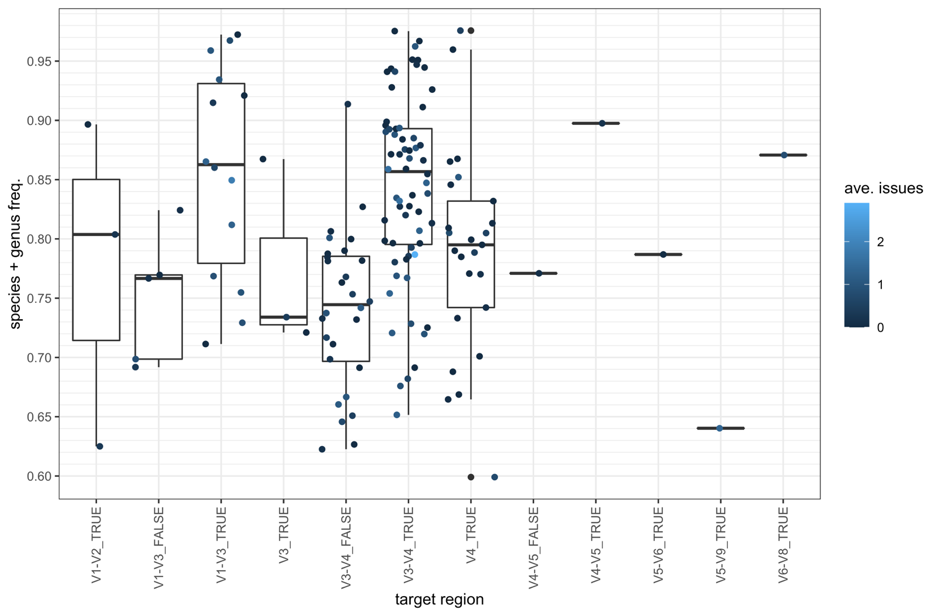

**Supplementary Table 5**. Median and 90° percentile values for the frequency of taxonomic assignment of ASVs at the species level in FoodMicrobionet studies from 34 to 180, by 16S gene target region. Only studies for which non-paired end sequences (IonTorrent or 454 platforms) or overlapping paired end sequence (Illumina platform) sequences are shown. n indicates the number of studies using a given target. Both the absolute frequency and the frequency weighted by sequence abundance are shown.

|  |  | Frequency of genus + species identification | | Frequency of genus + species identification, weighted by sequence abundance | |
| --- | --- | --- | --- | --- | --- |
| Target | n | median | 90° perc | median | 90° perc |
| V1-V2_TRUE | 3 | 0.167 | 0.189 | 0.266 | 0.260 |
| V1-V3_TRUE | 14 | 0.191 | 0.245 | 0.381 | 0.535 |
| V3_TRUE | 3 | 0.117 | 0.069 | 0.133 | 0.072 |
| V3-V4_TRUE | 64 | 0.205 | 0.113 | 0.301 | 0.453 |
| V4_TRUE | 25 | 0.137 | 0.072 | 0.209 | 0.220 |
| V4-V5_TRUE | 1 | 0.168 | 0.078 | 0.168 | 0.078 |
| V5-V6_TRUE | 1 | 0.069 | 0.004 | 0.069 | 0.004 |
| V5-V9_TRUE | 1 | 0.025 | 0.003 | 0.025 | 0.003 |
| V6-V8_TRUE | 1 | 0.313 | 0.106 | 0.313 | 0.106 |

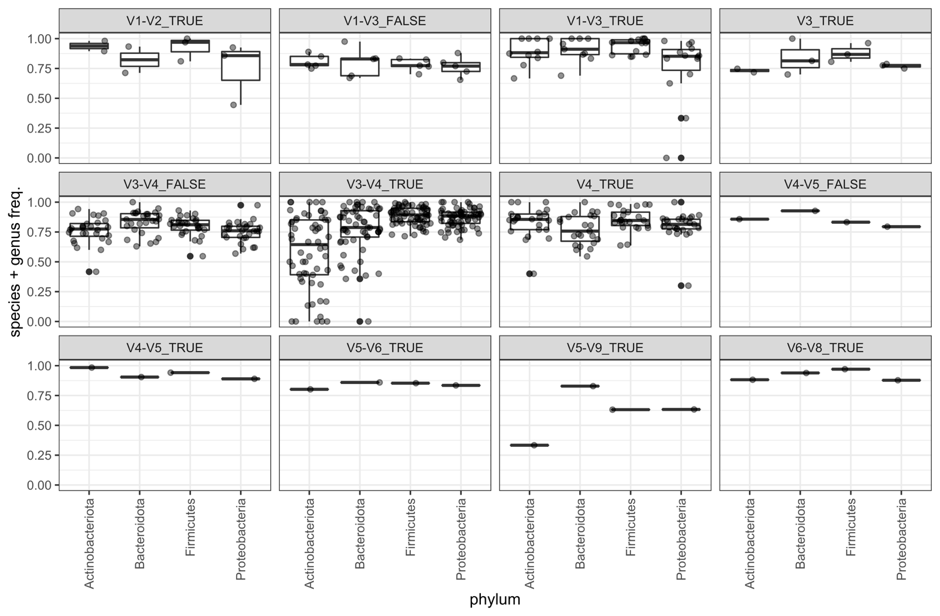

**Supplementary figure 4.** Box and jitter plots showing the unweighted (sequence relative abundance ignored) distribution of frequencies of taxonomic assignments at the genus level or below in FoodMicrobionet studies 34 to 180 for the four most abundant phyla.

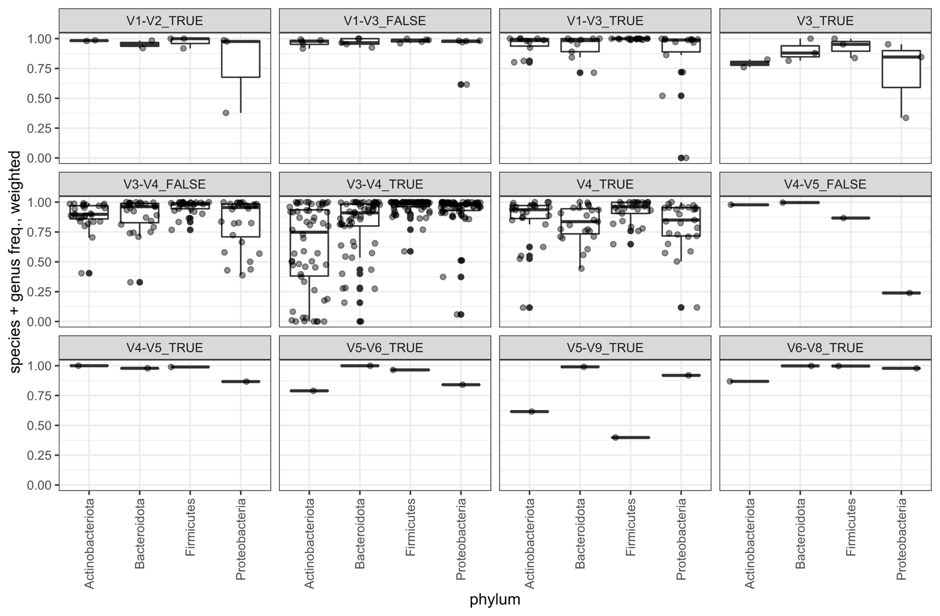

**Supplementary figure 5.** Box and jitter plots showing the weighted (by sequence relative abundance) distribution of frequencies of taxonomic assignments at the genus level or below in FoodMicrobionet studies 34 to 180 for the four most abundant phyla.

**Supplementary Table 6.** Data on the distribution of *Salmonella* and *Listeria in* food samples in FoodMicrobionet. The samples are divided in groups using the L1 of EFSA FoodEx2 classification. The number of samples for each group (n), the prevalence of a given genus in a sample group (as fraction of the samples in which the genus is found) and the median, median absolute deviation (mad) and maximum (max) abundance (%) are also shown.

|  |  |  |  | abundance (%) | | |
| --- | --- | --- | --- | --- | --- | --- |
| Genus | L1 | n | prev. | median | mad | max |
| *Listeria* | Animal and vegetable fats and oils and primary derivatives thereof | 53 | 0.8679 | 0.0057 | 0.0075 | 0.2113 |
|  | Fish, seafood, amphibians, reptiles and invertebrates | 461 | 0.0803 | 0.0018 | 0.0025 | 0.0201 |
|  | Legumes, nuts, oilseeds and spices | 49 | 0.1020 | 0.0001 | 0.0001 | 0.0028 |
|  | Meat and meat products | 1836 | 0.0321 | 0.0012 | 0.0015 | 0.1477 |
|  | Milk and dairy products | 4499 | 0.0080 | 0.0007 | 0.0010 | 0.0776 |
|  | Vegetables and vegetable products | 604 | 0.0017 | 0.0007 | 0.0000 | 0.0007 |
| *Salmonella* | Alcoholic beverages | 114 | 0.0702 | 0.0005 | 0.0003 | 0.0011 |
|  | Animal and vegetable fats and oils and primary derivatives thereof | 53 | 0.0566 | 0.0001 | 0.0000 | 0.0002 |
|  | Fish, seafood, amphibians, reptiles and invertebrates | 461 | 0.0130 | 0.0705 | 0.0360 | 0.1527 |
|  | Grains and grain-based products | 58 | 0.0517 | 0.0023 | 0.0017 | 0.0035 |
|  | Major isolated ingredients, additives, flavours, baking and processing aids | 280 | 0.0250 | 0.0095 | 0.0125 | 0.0538 |
|  | Meat and meat products | 1836 | 0.0926 | 0.0021 | 0.0018 | 0.5137 |
|  | Milk and dairy products | 4499 | 0.0127 | 0.0004 | 0.0004 | 0.0265 |
|  | Seasoning, sauces and condiments | 138 | 0.0072 | 0.0005 | 0.0000 | 0.0005 |
|  | Vegetables and vegetable products | 604 | 0.0430 | 0.0020 | 0.0024 | 0.0280 |

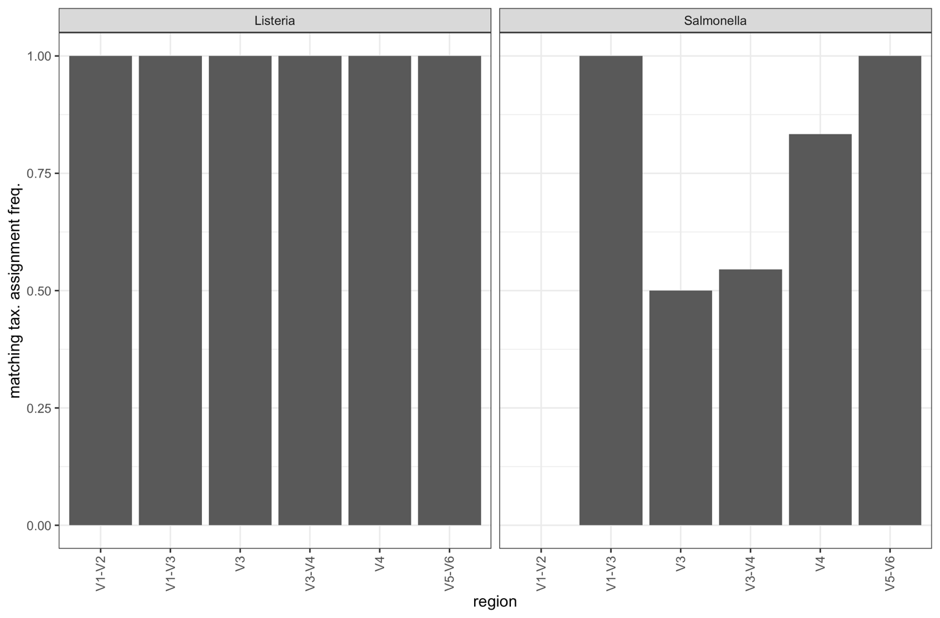

**Supplementary figure 6.** Frequencies of matching taxonomic assignments at the genus level with the SILVA v138.1 and RDP v18 trainset reference databases for ASVs assigned by SILVA to *Salmonella* or *Listeria*.

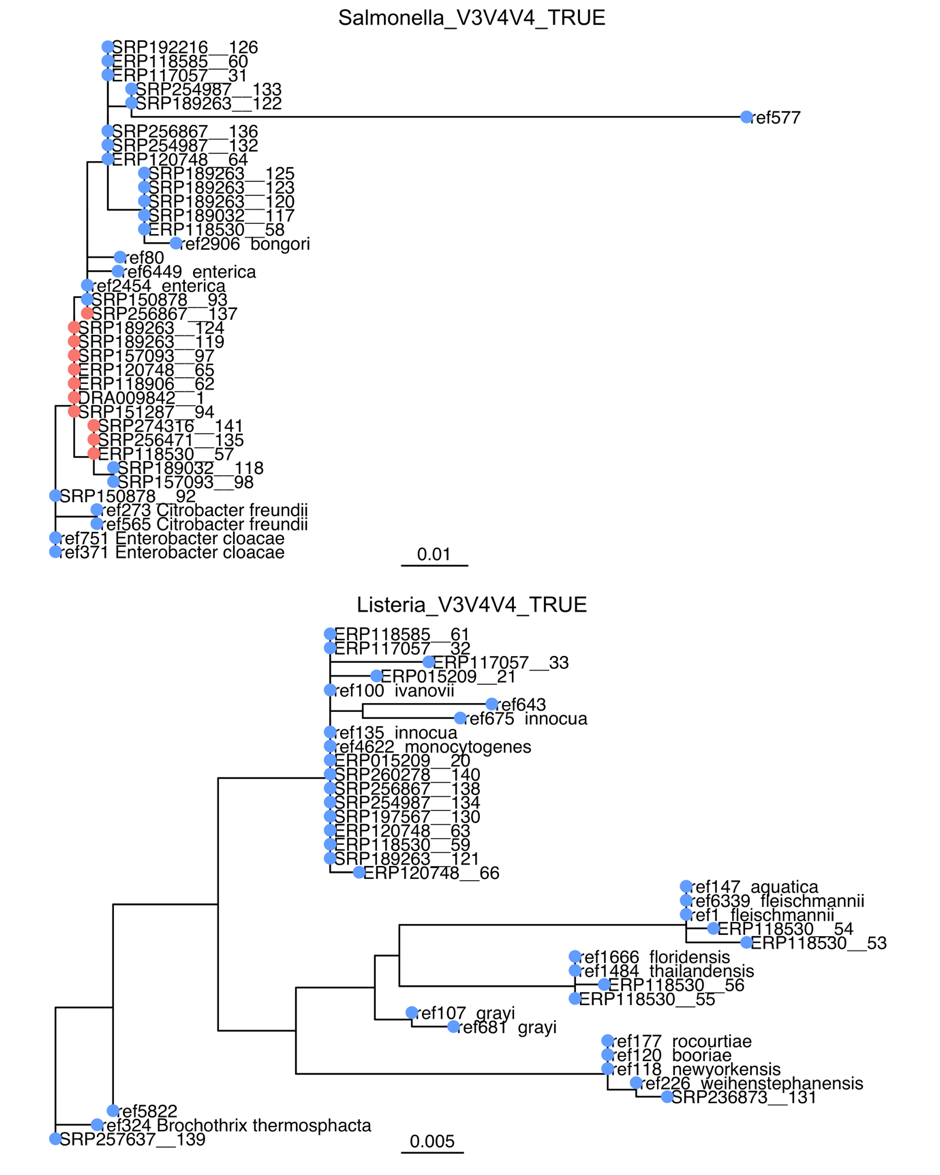

**Supplementary figure 7.** Maximum likelihood phylogenetic trees for amplicon sequence variants (ASVs) for overlapping paired end sequences for region V3-V4 and V4 (this means that alignment was performed for the V4 region only) of the 16S RNA gene identified as *Salmonella* or *Listeria*. ASVs are identified by the accession number of the study to which they belong and by a random progressive integer. Reference sequences extracted from SILVA v138.1 taxonomic reference database are also included. Colored dots indicate sequences for which taxonomic assignment with SILVA v138.1 and RDP trainset 18 matched (blue) or not (red).
